# Interrogation of noncoding schizophrenia risk variants using CRISPR-based functional genomics

**DOI:** 10.64898/2026.09.03.749158

**Authors:** Marisa C. Hamilton, Julia W. Riley, Alexander C. Nelson, Boxun Li, Sophie F. Dornbaum, Alexias Safi, Xiekui Cui, Ian R. Jones, Aaron A. Coley, Kevin T. Hagy, Ruhi Rai, Alejandro Barrera, Andrew S. Allen, Yin Shen, Richard I. Sherwood, Michael I. Love, Patrick F. Sullivan, Charles A. Gersbach, Gregory E. Crawford

## Abstract

Schizophrenia (SCZ) is a highly heritable complex disorder influenced by coding and noncoding genetic variation. Its genetic causes, particularly those involving noncoding variation, are largely unknown. High-throughput CRISPR screens enable dissection of disease-associated loci and identification of noncoding regulatory elements and variants that modulate gene expression. We screened SCZ GWAS loci linked to genes that are also associated in whole-exome sequencing studies to identify regulatory elements and variants impacting expression of disease-relevant genes. We used CRISPRi paired with HCR-FlowFISH to epigenetically silence 333 putative regulatory elements and measure the downstream effects on gene expression of causal SCZ genes, *FAM120A*, *SV2A*, and *STAG1*, in iPSCs and iPSC-derived neurons (iNeurons). We identified 78 regulatory elements that significantly alter expression of a SCZ gene, including noncoding enhancers/silencers as well as promoters of genes and lncRNAs. Pooled prime editing screens interrogated noncoding variant influence on gene expression for SCZ-associated variants and uncharacterized common variants from diverse population studies. We find that a common variant in the promoter of *SV2A*, rs112851681:A>G (MAF = 3.56%, 1000 Genomes) enhances transcriptional activity in iPSCs and iNeurons. These findings show distinct noncoding mechanisms that map within GWAS signals, and provide a path forward for interrogating noncoding regulatory elements and variants in disease loci.

## Introduction

Schizophrenia (SCZ) is a chronic neuropsychiatric disorder with a lifetime risk of ∼1%^1^ and is associated with increased morbidity and mortality. Antipsychotics treat only a portion of symptoms in some individuals and often cause clinically-important side-effects. Approximately 30% of patients are categorized as treatment resistant for unknown reasons^2,3^, supporting a crucial need for more effective therapies. A major roadblock in the discovery of novel therapeutics to treat SCZ is the gap in understanding of the biological mechanisms driving this disorder. Recent advances in the fields of genetics and genomics have allowed for substantial progress in identifying regions of the genome, individual genes, and cell-types involved in SCZ^4–6^.

Genome-wide association studies (GWAS) have uncovered a contribution of common variants of small effects to the genetic architecture of SCZ^5^. A recent GWAS identified >22,000 associated variants in 287 distinct loci where common variation is associated with SCZ risk^5^. Additionally, whole-exome sequencing (WES) of SCZ cases identified 32 genes (FDR <0.05) containing rare variants that confer an increased risk of SCZ^6^. There is an enrichment for overlap between the loci identified in the SCZ GWAS and genes found in the SCZ WES study^5^, indicating that common and rare variant associations are converging on a subset of the same genes. This pattern of converging GWAS and WES signals has also been observed in other psychiatric and non-psychiatric traits with both polygenic and monogenic contributions to risk, such as autism spectrum disorder, kidney disease, body mass index and coronary artery disease^7–10^. The majority of common disease-associated GWAS variants map to noncoding regions, making it difficult to interpret which gene(s) they impact. However, genes identified through WES studies that lie near or within GWAS signals offer a disease-relevant set of genes to investigate nearby regulatory variants that contribute to SCZ risk.

SCZ GWAS variants are enriched near genes expressed in neurons of the central nervous system, making neurons a prioritized cell-type in SCZ^5,11^. With current technologies, primary human neurons are not suitable for high-throughput screens. Previous studies have established rapid and homogenous differentiation of induced pluripotent stem cells (iPSCs) into glutamatergic neurons (iNeurons) through overexpression of the murine ortholog of *NEUROG2*, *Ngn2*^12,13^. These neurons are amenable to high-throughput CRISPR screening to investigate neuronal differentiation and disease^5,14^ and have been pivotal in the study of genes and pathways underlying risk for neuropsychiatric disorders^15–17^.

Advancements in the CRISPR toolkit have enabled targeted genetic and epigenetic manipulation of the genome, facilitating high-throughput interrogation of disease-associated loci in iPSC-derived cellular models^18^. Using a nuclease-deactivated Cas9 (dCas9) fused to repressive domains (CRISPRi)^19–22^ allows for large-scale characterization of putative regulatory elements (pREs) in disease-associated loci^23–25^. Multi-gene transcriptomic readouts combining CRISPRi with single-cell RNA-sequencing has allowed for unbiased linking of pREs to nearby genes^26^, which has significant utility in loci where there are a large number of possible gene targets. However, these approaches currently have limited sensitivity for identifying REs with small effect sizes, struggle to detect lowly expressed genes, and do not readily enable functional dissection of individual variants at scale^27–29^. Moreover, while unbiased screens can link REs to their target genes, interpreting the function of genetic variants that map within these REs in disease risk remains challenging when the target genes have an unknown role in disease pathology.

In loci with a distinct disease-relevant gene target, such as those overlapping or linked to SCZ WES genes, hybridization chain reaction fluorescence *in situ* hybridization coupled with flow cytometry (HCR–FlowFISH^30^) offers a scalable and highly sensitive gene expression readout for mapping functional REs and variants. While CRISPRi HCR-FlowFISH can successfully link REs to target genes^30^, additional CRISPR tools can be paired with HCR-FlowFISH to investigate variant effects on gene expression. Prime editing (PE) utilizes a Cas9 nickase and reverse transcription (RT) enzyme along with a prime editing gRNA (pegRNA) to induce precise edits^31–33^, including indels, and has been combined successfully with HCR-FlowFISH to characterize synthetic noncoding variants in high-throughput^34^. Nonetheless, PE paired with HCR-FlowFISH has yet to be applied to characterize common noncoding variants in GWAS loci.

Because SCZ GWAS variants predominantly map to noncoding regions, we systematically interrogated REs and variants within GWAS loci using pooled CRISPR screens in iPSCs and iNeurons. To identify REs and variants influencing expression of a disease-relevant gene target, we investigated genes prioritized by SCZ WES that are also linked to a SCZ GWAS locus. We characterized three SCZ GWAS loci through pooled CRISPRi perturbation of 333 total pREs, identifying 78 REs that control expression of known SCZ genes (*FAM120A*, *STAG1*, and *SV2A*). We next used a combination of pooled PE screens, targeted PE, reporter assays, and computational predictions to identify noncoding variants that influence RE activity. Overall, our systematic approach to dissect SCZ-associated loci reveals noncoding REs and variants that modulate known SCZ genes. This screening strategy can be applied to the study of other polygenic disorders to better understand how REs and noncoding variants contribute to disease risk.

## Results

### Identification of SCZ GWAS putative gene targets using WES

To nominate putative gene targets for CRISPR-based interrogation of SCZ GWAS loci, we identified GWAS^5^ regions that map within ±1 Mb of genes implicated in SCZ by WES^6^ (**Figure 1A, Table S1**). SCZ WES genes are significantly enriched in SCZ GWAS loci^5^, and the assumption is that such genes are involved in SCZ in at least two ways: rare coding variants of large effect size and common noncoding variants of small effect size that modulate gene expression. Consistent with this assumption, using the Missense badness, PolyPhen-2, and Constraint (MPC) score, we find that eight WES genes showed stronger effects for highly pathogenic class I coding variants (i.e., MPC > 3) compared to more moderately pathogenic class II coding variants (MPC 2-3)^6^ (**Figure 1B**). This suggests that common noncoding variants with small effects on the same genes could also influence SCZ risk. Five GWAS loci overlapped WES-prioritized genes, and five others contained either eHi-C (fetal or adult cortex^35^) or GTEx expression quantitative trait locus (eQTL) links to the WES-implicated proximal gene, supporting *cis*-regulatory interactions (**Figure S1A, Table S1**). The genomic context was variable, with 5-192 protein-coding genes and 1-138 fine-mapped SNPs^5^ per locus, indicating a substantial number of putative causal genes and variants for many loci (**Figure S1B-C**). The majority of GWAS fine-mapped SNPs mapped to noncoding regions, suggesting that common disease-associated variants act primarily through gene regulatory mechanisms, as found for other common disorders^36,37^ (**Figure S1D**).

**Figure 1:**
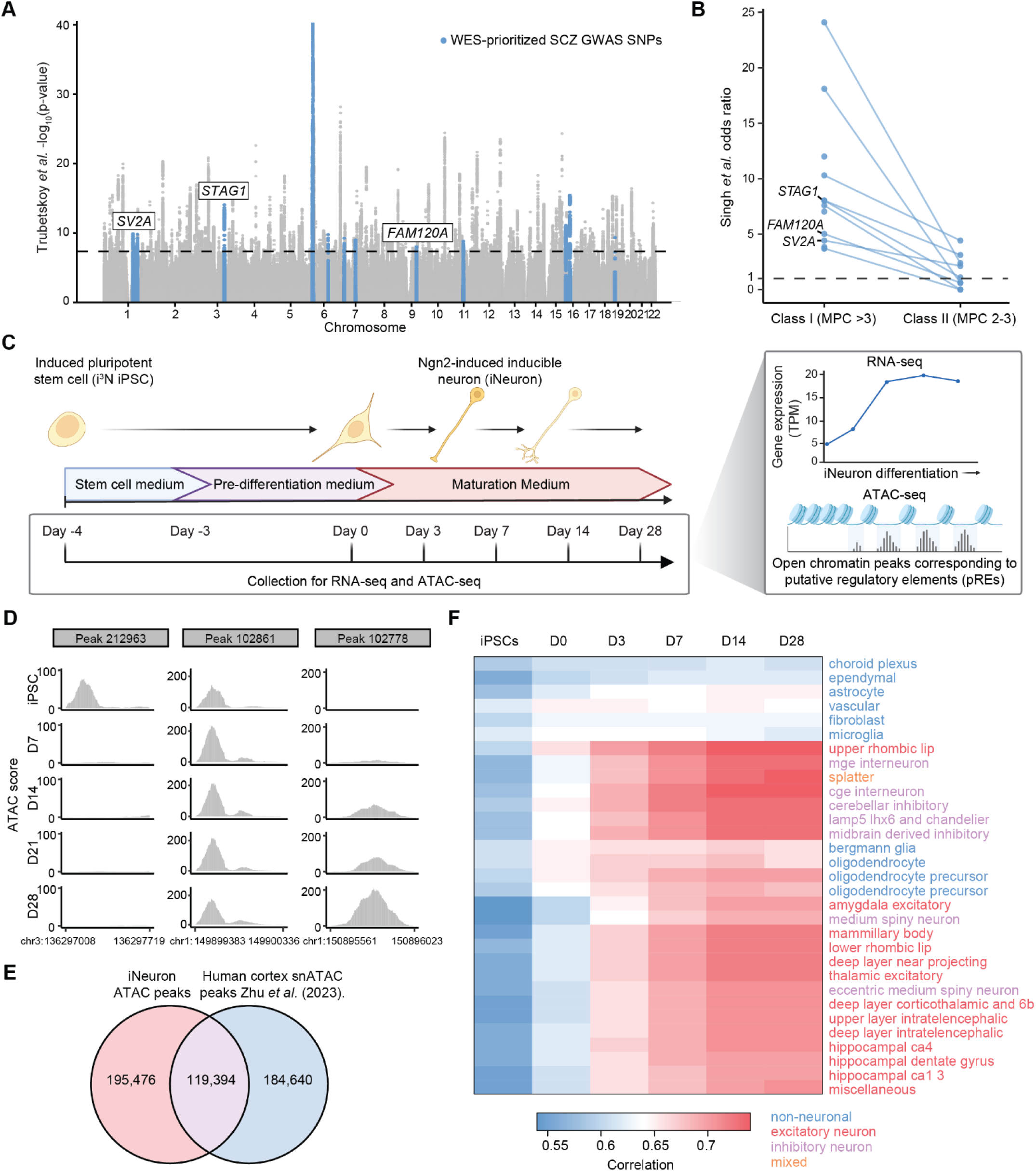
SCZ WES genes converge with SCZ GWAS signal at 13 genomic loci. A) Manhattan plot depicting Trubetskoy *et al.* SCZ GWAS data for all imputed SNPs. The dashed line indicates genome-wide significance threshold. Imputed SNPs in SCZ-associated GWAS loci ± 1 Mb of a SCZ-WES gene are highlighted in blue. GWAS loci prioritized for CRISPRi screening are labeled by the SCZ WES gene directly overlapping or proximal to the locus. B) Reported odds ratios of SCZ WES genes ± 1 Mb of a SCZ GWAS locus, separated into two classes of MPC pathogenicity score. SCZ WES genes directly overlapping or proximal to a SCZ GWAS locus prioritized for CRISPRi screening are labeled. C) Schematic depicting four week iNeuron differentiation with ATAC-seq and RNA-seq collected at D-4 (iPSCs), D0, D7, D14, D21, and D28. ATAC-seq was used to identify pREs and RNA-seq was used to determine gene expression levels. D) ATAC scores across the iNeuron differentiation for three representative consensus peaks at five representative timepoints. E) Venn diagram showing the overlap between the union set of ATAC-seq peaks identified in iNeurons (red) and the union set of snATAC-seq peaks from six developmental timepoints of the human cortex (blue)^39^. F) Spearman correlations (ρ) of gene expression of protein-coding genes in the iPSC to iNeuron differentiation with gene expression in the 31 superclusters in the Human Brain Atlas (single-nucleus RNA-seq in adult brains)^40^.

To study gene regulatory mechanisms using a tractable cell model system, we use an iPSC dCas9^KRAB^ line containing a doxycycline-inducible *Ngn2* transgene (i^3^N iPSCs^38^) and differentiate it across six excitatory neuron stages: Day −4 (iPSCs), D0 (neural progenitor-like), and D3/7/14/28 iNeurons (glutamatergic neurons) (D = “day in maturation medium”) (**Figure 1C**). To confirm this is a disease-relevant cell type for modeling gene regulation within these loci, we generated RNA-seq and ATAC-seq datasets for these cells across all time points (**Figure 1C**). Across these stages, we detected 13,569 protein-coding genes with mean TPM > 1 and a union set of 314,870 ATAC-peaks **(Figure 1D, Tables S2-3)**. As expected, gene expression in iPS and D0 cells had higher correlations with each other, while D3-D28 cells showed higher correlations **(Figure S1E).**

We overlapped the union set of iNeuron ATAC peaks with a union set of snATAC-seq peaks from six developmental timepoints of the human cortex^39^ and found that 40% of accessible chromatin regions from human brain samples were represented during iPSC-to-iNeuron differentiation (**Figure 1E**). As the brain snATAC-seq dataset spans all cortical cell types rather than neurons alone, this comparison likely underestimates the true overlap with specifically the neuronal lineage. We then compared the gene expression profiles of the iPSCs and D0-28 iNeurons to RNA-seq data from Human Brain Atlas^40^ (31 superclusters, **Figure 1F**). Spearman correlations were the lowest between iPSCs and neuronal subtypes. The correlations with neuronal subtypes increased with differentiation time, with the highest correlations between D14 and D28 iNeurons. There were correlations ≥ 0.7 with all 13 excitatory neuron superclusters and 6 of 7 inhibitory neuron superclusters. Only one non-neuronal subtype had a correlation exceeding 0.70 **(Figure 1F)**.

Of the 13 SCZ WES genes ± 1 Mb of a SCZ GWAS locus, 12 were expressed (TPM ≥ 5) in iPSCs or D0-28 iNeurons, indicating that the majority of these disease-relevant genes are transcriptionally active in these cell models and enabling investigation of gene regulatory mechanisms using CRISPRi. (**Figure S1F, Table S1**). ATAC-seq identified a median of 69 open chromatin peaks per locus, which we used to define pREs **(Figure S1F**). In addition, 12 WES-prioritized loci colocalized with iPSC chromatin accessibility QTL (caQTL), histone acetylation QTL (haQTL), or eQTL signals, with 11 loci colocalizing with early development (EDev)-specific or EDev-shared eQTLS^41^ (**Figure S1G**). This indicates that genetic variation in these SCZ-associated loci actively modulates gene regulation in iPSCs and at early developmental states. Taken together, these results suggest iPSCs and iNeurons are relevant models for dissecting WES-prioritized SCZ-associated loci and support that gene regulation during early fetal development may influence SCZ risk.

### CRISPRi screening identifies noncoding regulators of WES-identified SCZ genes

CRISPRi HCR-FlowFISH can robustly assay pREs for regulatory function and link REs to gene targets^30^ **(Figure 2A)**. We proceeded with CRISPRi HCR-FlowFISH screening in both iPSCs and iNeurons for three WES-prioritized loci, which were chosen based on direct overlap or proximity of a GWAS locus to the WES gene (*STAG1*, *FAM120A*, *SV2A*), existence of eHi-C and/or eQTL connections, and expression levels in iPSCs and iNeurons **(Figure S1F, Table S1)**. We designed three gRNA libraries targeting pREs identified from the union set of ATAC-peaks across the iPSC-to-D28 differentiation time course. We expanded the boundaries of these GWAS loci by 10% to capture pREs that may harbor functional variants with strong effects but are in low LD with the index SNP. The three gRNA libraries each included ∼2,000 gRNAs and 200 non-targeting gRNA controls, with a total of 333 targeted pREs (mean of 17 gRNAs/pRE) **(Tables S4-6)**. We transduced each of the three pRE targeting libraries into iPSCs and used HCR-FlowFISH to cytometrically sort cells into bins based on low (0-15 percentile) or high (85-100 percentile) target gene expression, normalizing for variability in size and permeability using the housekeeping gene *TBP* (**Figure 2A, Figure S2A**). We additionally screened the *SV2A* pRE targeting library in Day 7 iNeurons given that *SV2A* is 4.5X more highly expressed in iNeurons (**Figure S1F**). We sequenced gRNAs from each bin, observing a strong correlation in gRNA counts across replicates for both iPSCs and iNeurons, indicating high reproducibility **(Figure S2B-E**).

**Figure 2:**
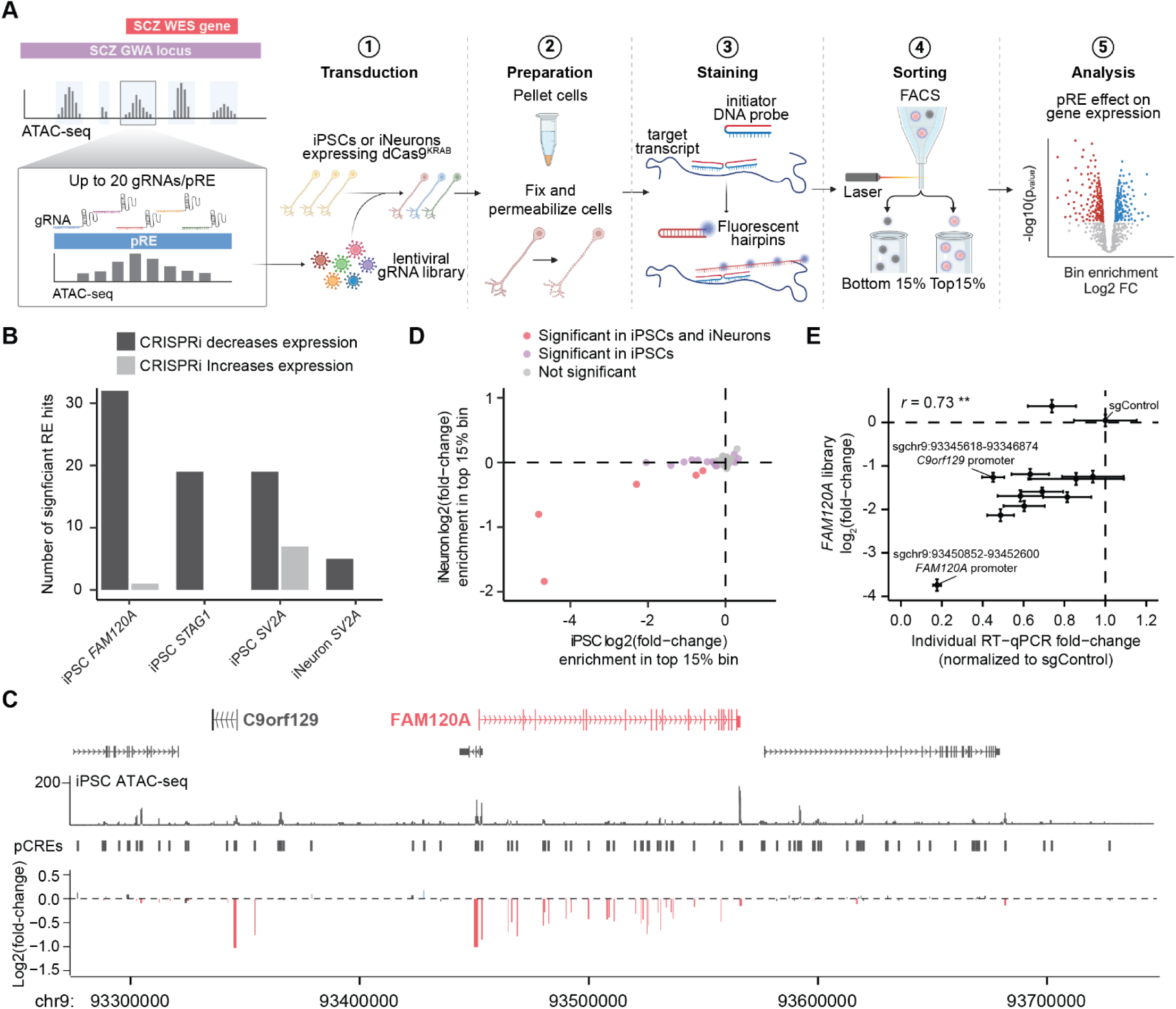
CRISPRi screens identify REs in SCZ GWAS loci that modulate expression of known SCZ genes. A) Schematic of HCR-FlowFISH CRISPRi regulatory element screening in iPSCs and iNeurons. B) Number of significant (adjusted p-value <= 0.05) regulatory element hits in each iPSC or iNeuron screen. C) Locus view displaying the *FAM120A* GWAS locus. From top to bottom: GENCODE V48 genes and predicted genes; i^3^N iPSC ATAC-seq scores; pREs identified through called ATAC-seq peaks; pRE level log_2_(fold-change) in enrichment in top vs bottom 15% bins after *FAM120A* HCR-FlowFISH and FACS sorting of i^3^N iPSCs transduced with the FAM120A locus pRE targeting gRNA library. D) Comparison between pRE log_2_(fold-changes) from *SV2A* CRISPRi HCR-FlowFISH screening in iPSCs and iNeurons. E) Pearson correlation between expression of *FAM120A* in an individual gRNA validation using RT-qPCR and the *FAM120A* pRE library gRNA log_2_(fold-change) in HCR-FlowFISH screening of iPSCs. n=3-4. (∗p < 0.05, ∗∗p ≤ 0.01, ∗∗∗p ≤ 0.001).

Analysis with MAGeCK^42^ identified 78 REs that significantly modulate expression of a known SCZ WES gene in iPSCs or iNeurons (70 decrease expression, 8 increase expression at p_adj_ ≤ 0.05; **Figure 2B-C, Figure S3, Figure S4, Tables S7-9).** For both iPSCs and iNeurons, the respective gene promoters were the top hits **(Figure 2C, Figure S3A-G).** We observed a partial correlation in RE effect size in our iPSC and iNeuron screens in the *SV2A* locus, suggesting there are both shared and unique mechanisms for gene regulation between these two cell types (**Figure 2D**). To further validate the accuracy of our screens in both cell types, we performed RT-qPCR for 12 gRNAs in iPSCs and 10 gRNAs in iNeurons, observing a strong correlation between the fold-change in enrichment observed in the screens and the fold-change in gene expression identified through individual RT-qPCR validation **(Figure 2E, Figure S4A).**

### CRISPRi-identified REs encompass both putative cis- and trans-mechanisms of gene regulation

We next characterized RE hits identified in iPSC and/or iNeurons. The CRISPRi-identified REs spanned both genic and intergenic regions, with the majority of REs mapping in introns belonging to the target gene (for *FAM120A* and *STAG1*) or promoters of other genes within the locus (for *SV2A*) **(Figure S5A)**. The median distance of each RE to the transcription start site of the target gene was 84 kb, with the total range of distances spanning from 0 kb (promoter) to 1.28 Mb **(Figure S5B).** Of the 69 REs located more than 5 kb from a TSS of a WES gene, seven were supported by eHi-C chromatin loops in iPSCs, iNeurons, fetal cortex, and/or adult cortex^35^ **(Figure S5C)**. This supports that at least a subset of our CRISPRi-identified REs influence gene expression through *cis*-acting mechanisms mediated by chromatin looping. The remaining REs not supported by eHi-C chromatin loops may reflect weaker *cis*-interactions that are not readily detectable by eHi-C, *cis*-interactions present in our cell lines that are not present in the eHi-C assayed cell lines, or *trans*-interactions.

To elucidate possible mechanisms of these GWAS loci and prioritize common SCZ-associated variants, we combined our CRISPRi RE results with other lines of evidence (**Table S11**). In one example, we identified a RE ∼300 kb upstream of *SV2A* that is supported by a fetal cortex eHi-C chromatin interaction^35^ (**Figure 3A**). This RE contained a FINEMAP SCZ SNP (95% credible set)^5^, rs72694957:T, which was also identified as an eQTL in GTEx that significantly decreases *SV2A* expression in 10 GTEx tissues, including cultured fibroblasts **(Figure 3B)**. While this SNP was not a significant eQTL in any bulk brain tissues, we found that this SNP resulted in a trending decrease in *SV2A* expression in the frontal cortex **(Figure 3B)**, likely due to either smaller sample sizes for brain tissues in GTEx or GTEx using bulk RNA-seq^43^. A recent study using a massively parallel reporter assay (MPRA) in primary human neural progenitors reported decreased regulatory activity for rs72694957:T, further supporting that this SNP can influence transcription^44^. MPRA is a powerful tool to identify allelic regulatory effects in isolation, but does not directly connect variants to target genes. Our CRISPRi screen provides evidence directly linking rs72694957:T to *SV2A* expression, providing additional support for this variant contributing to SCZ risk.

**Figure 3:**
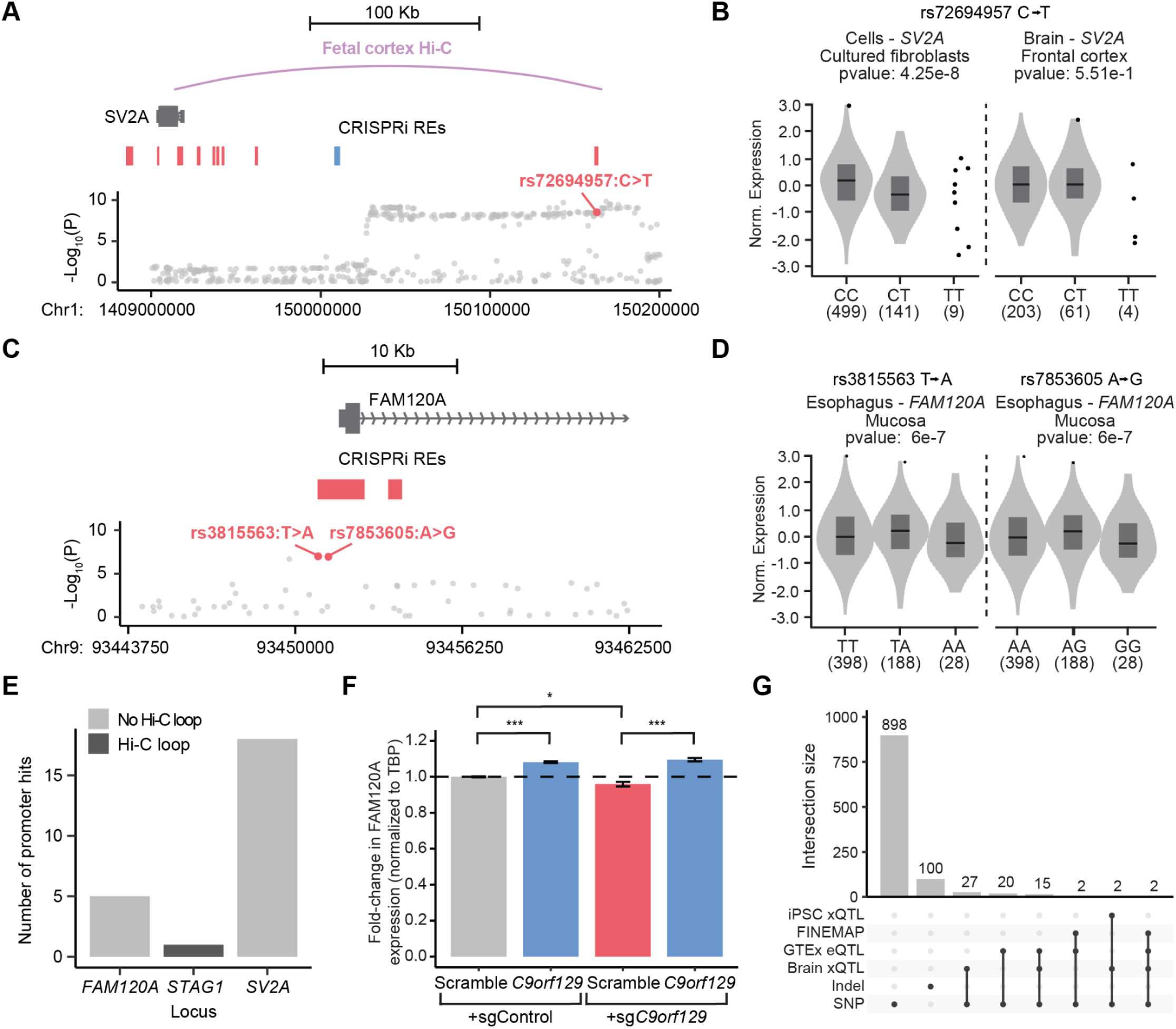
CRISPRi screens prioritize noncoding variants in SCZ-associated loci. A) Locus view displaying a fetal cortex eHi-C chromatin loop between a CRISPRi-identified RE and the promoter of *SV2A.* Blue = CRISPRi increased expression of *SV2A*, red = CRISPRi decreased expression of *SV2A*. The bottom row depicts imputed SCZ GWAS SNPs and their corresponding -log_10_(p-values) from *Trubetskoy* et al. A finemapped SCZ SNP directly overlapping the eHi-C linked RE, rs72694957, is labeled in red. B) Violin plots of normalized *SV2A* expression across rs72694957 genotypes (CC, CT, TT) in cultured fibroblasts (left) and frontal cortex (right) from GTEx. C) Locus view displaying the promoter of *FAM120A.* Red = CRISPRi decreased expression of *FAM120A*. The bottom row depicts imputed SCZ GWAS SNPs and their corresponding -log_10_(p-values) from *Trubetskoy* et al. Two finemapped SCZ SNPs directly overlapping the *FAM120A* promoter, rs3815563 and rs7853605, are labeled in red. D) Violin plots of normalized *FAM120* expression across rs3815563 (left) and rs7853605 (right) genotypes (TT, TA, AA and AA, AG, GG) in esophagus mucosa from GTEx. E) Bar plot showing the number of RE promoter hits for *FAM120A*, *STAG1*, and *SV2A* with or without supporting eHi-C chromatin interactions from iPSCs, iNeurons, fetal cortex, or adult cortex. REs overlapping *FAM120A*, *STAG1*, or *SV2A* promoters were excluded. F) Normalized fold-change in *FAM120A* expression following overexpression of *C9orf129* lncRNA compared to scramble control in i^3^N iPSC expressing sg*C9orf129* or sgControl as measured by HCR-FlowFISH. n=3. Two-tailed t test. G) Upset plot showing the overlap of common variants with functional signals including QTL datasets^45,46,41^ and SCZ FINEMAP variants (95% credible set)^5^. (∗p < 0.05, ∗∗p ≤ 0.01, ∗∗∗p ≤ 0.001).

In addition to putatively functional distal variants, we also identified 157 common variants (All dbSNP (155), minor allele frequency (MAF) ≥ 1% in any project) within ATAC-seq peaks overlapping the TSSs of *FAM120A*, *STAG1*, and *SV2A* (**Figure S5D**). Notably, two of these variants are SCZ FINEMAP SNPs located within the *FAM120A* promoter, rs3815563 and rs7853605 (**Figure 3C**). Interestingly, both variants were associated with a significant increase in *FAM120A* expression in esophageal mucosa in GTEx **(Figure 3D)**. Mechanistically, common promoter variants may mediate these effects by disrupting or improving TF binding motifs or chromatin accessibility, and thereby altering transcription initiation. Given that rs3815563 and rs7853605 are in strong LD, further experimental validation will be required to resolve their individual contributions to *FAM120A* regulation and SCZ risk.

Most REs lacked support from chromatin looping data, which may reflect the limited sensitivity of eHi-C or indicate that some REs act through *trans*-mechanisms. Across the three SCZ-associated loci, our CRISPRi screens identified 24 RE hits mapping to promoter regions of nearby genes **(Figure 3E)**, yet only 1 of these REs had an eHi-C interaction with a WES gene promoter. This suggests potential *trans*-regulatory effects for a subset of REs. As one example, a RE mapped to the promoter of a pseudogene/lncRNA, *C9orf129,* and targeting this region by CRISPRi resulted in an ∼50% reduction in *FAM120A* expression (∼100 Kb away) in iPSCs (**Figure 2E**). *C9orf129* is more highly expressed in brain tissue compared to all other GTEx tissues **(Figure S5E)** and shows high co-expression (*R* = 0.92) with *FAM120A* in neurons profiled in a single-cell atlas of the developing brain **(Figure S5F),** indicating a putative role of this lncRNA promoter (*cis*) and/or lncRNA (*trans*) in influencing expression of *FAM120A* in the developing brain. We assessed whether *C9orf129* could modulate *FAM120A* expression independently of its genomic context by exogenously overexpressing *C9orf129* in iPSCs with or without endogenous knockdown of *C9orf129* (**Figure S5G).** Overexpression led to a significant increase in *FAM120A* transcript levels by HCR-FlowFISH in both the endogenous knockdown and endogenous wild-type *C9orf129* conditions (**Figure 3F**) and a non-significant but trending increase in *FAM120A* transcript levels by RT-qPCR **(Figure S5H)**. Complete rescue of *FAM120A* expression by exogenous *C9orf129* despite *C9orf129* promoter CRISPRi, as measured by HCR-FlowFISH, suggests a *trans*-acting mechanism mediated by *C9orf129* lncRNAs.

### CRISPRi RE screening prioritizes common variants in SCZ-associated loci

We examined common genetic variants (MAF ≥ 1% in All dbSNP build 155) located within CRISPRi-identified REs. Many of these variants are not represented in GWAS and QTL analyses due to sample bias in the genetic backgrounds of participants/cell lines and, in some cases, due to imputation limitations (e.g., indels). In total, we identified 1,066 common variants mapping within our 78 CRISPRi identified REs, including 100 indels **(Figure S5I)**. To assess potential regulatory function, we overlapped these variants with FINEMAP-prioritized SCZ SNPs^5^ as well as a number of QTL datasets, including GTEx eQTLs, adult cortex eQTLs, haQTLs, and mQTLs^45^, fetal cortex eQTLs^46^, and iPSC eQTLs, caQTLs and haQTLs^41^. This analysis identified 68 SNPs that overlap with at least one of these functional signals **(Figure 3G, Table S11)**. We observed no overlap in functional signals with indels due to their underrepresentation in association and QTL datasets.

### A prime editing (PE) reporter quantifies pegRNA endogenous editing efficiencies

To enable large-scale investigation of variants, including indels, we optimized a PE screening platform in iPSCs. PE is a versatile and precise genome-editing technique that allows for the study of genetic variants in their native genomic contexts^47^. However, low editing efficiencies and variation of editing efficiencies across pegRNAs have limited the accuracy of high-throughput PE screens^48^. To overcome this limitation, we utilized a PE reporter that couples each pegRNA with a synthetic target sequence on the same construct^49–51^ (**Figure 4A**). This vector expresses an IGFpm1-NFATC2IPp1-PE2 prime editor^52^ fused via a linker to the N-terminal domain of La protein (encoded by *SSB*) and a nuclear localization signal^50,53^ (IN-PE2-SSB). Given that measuring endogenous editing efficiencies for pegRNA libraries targeting hundreds of distinct regions is technically challenging, editing at the paired reporter site provides a scalable and quantitative proxy for pegRNA activity. This allows for the scaling of effect sizes in pooled PE screens, known as ‘activity-normalization^37,50,51^.

**Figure 4:**
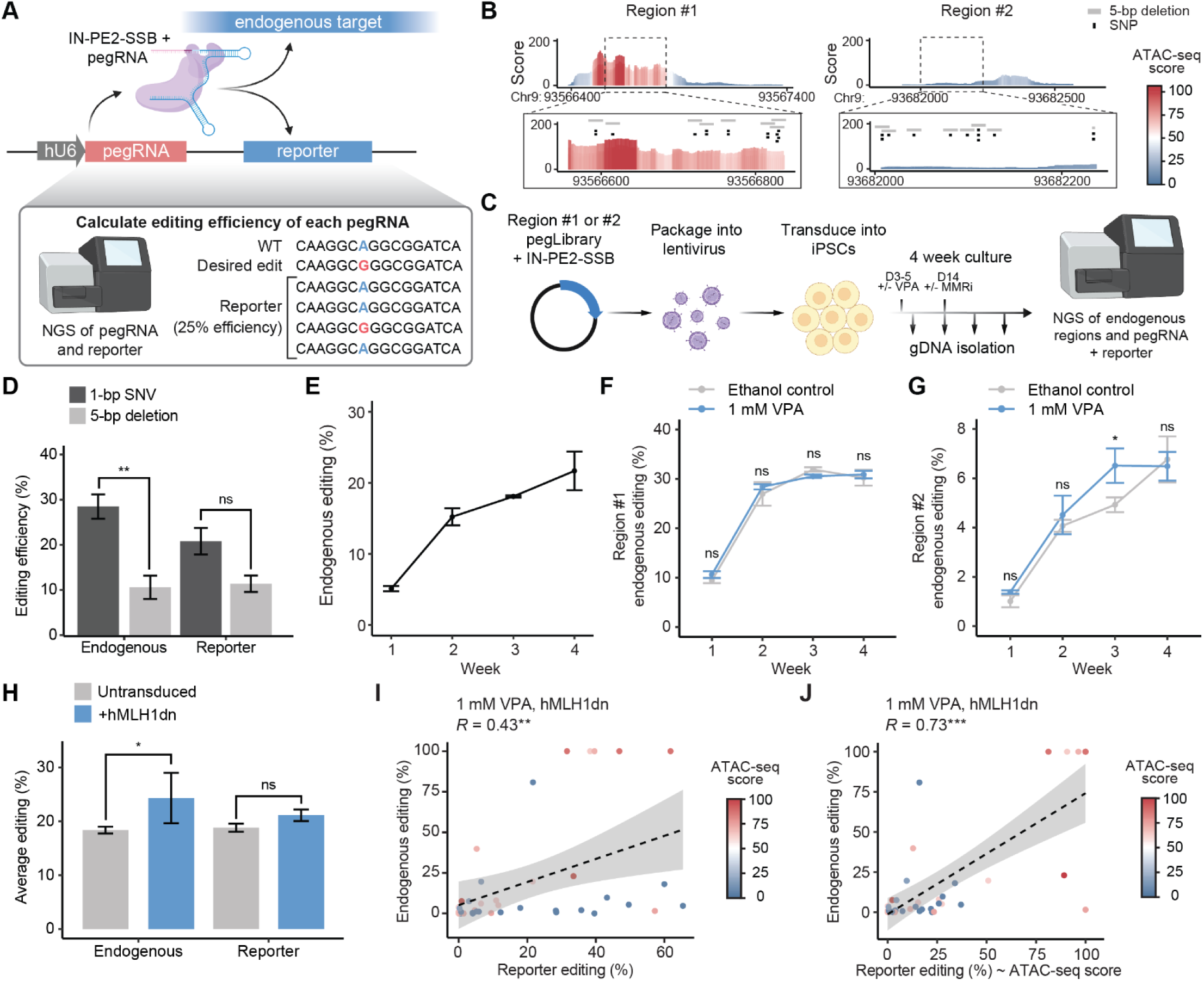
A prime editing reporter quantifies pegRNA endogenous editing efficiencies. A) Schematic of the pegRNA library reporter construct. The construct contains a pegRNA along with a pseudo-target reporter sequence that harbors the corresponding endogenous target site. pegRNAs edit at both the endogenous and pseudo-target sites. The pegRNA and reporter sequences are analyzed using NGS to obtain pegRNA identity and reporter editing efficiency. B) ATAC-seq tracks showing chromatin accessibility at the highly accessible locus (region #1) and the lowly accessible locus (region #2). SNP and 5-bp deletion pegRNA target sites are indicated. C) Experimental workflow. pegRNAs libraries targeting region #1 or region #2 were cloned into the reporter construct, packaged into lentivirus, transduced into iPSCs, cultured for four weeks, and both endogenous and reporter editing were quantified by NGS at weekly timepoints. D) Endogenous and reporter editing efficiencies for SNPs and 5-bp deletions at region #1 and region #2. n=3. Two-tailed t test. E) Average endogenous editing efficiency for all pegRNAs targeting region #1 and region #2 throughout a four-week time course. Average endogenous editing efficiency for all pegRNAs targeting (F) region #1 and (G) region #2 throughout a four-week time course with VPA or control treatment. Paired t test. H) Endogenous and reporter editing efficiencies at four weeks post-transduction of the pegRNA libraries with VPA treatment and with or without hMLH1dn lentiviral transduction. Paired t test. I) Pearson correlation between endogenous and reporter editing efficiency for all pegRNAs VPA and hMLH1dn treatment, measured at four weeks post-transduction. Each pegRNA is colored based on its average ATAC-score within a 50-bp window of the targeted variant. J) Pearson correlation of predicted (reporter editing (%) ∼ ATAC-seq score) versus observed endogenous editing efficiencies for pegRNAs following VPA and hMLH1dn treatment at four weeks post-transduction. (∗p < 0.05, ∗∗p ≤ 0.01, ∗∗∗p ≤ 0.001).

To validate the accuracy of the reporter construct in iPSCs, we transduced two pools of 20 and 22 pegRNAs, respectively, to install SNPs and 5-bp deletions across two genomic loci that had high (region #1) or low (region #2) chromatin accessibility **(Figure 4B, Tables S12-13)**. We sequenced both the endogenous and reporter sites over a four-week time course to assess editing outcomes over time (**Figure 4C**). Library diversity remained consistent throughout the four-week time course for both libraries, indicating a four week culture time does not cause a significant loss of diversity for these pegRNA pools (**Figure S6A-B**). Recombination rates (rate of chimeric pegRNAs as evidenced by spacer-barcode mismatches) were relatively consistent across libraries and throughout the timecourse, averaging 27.5% for region #1 pegRNAs and 24.8% for region #2 pegRNAs (**Figure S6C)**. While these recombination rates decrease signal-to-noise in PE screens, they are in line with previous reports for barcoded gRNA libraries^54,55^.

We observed that 5 bp deletions exhibited lower endogenous and reporter editing efficiencies compared to SNPs **(Figure 4D)**, consistent with prior reports for prime editors^56^. Both SNPs and 5 bp deletions showed increasing endogenous (**Figure 4E**) and reporter editing (**Figure S6D)** efficiencies over the four-week time course. PRIDICT2.0^56^ pegRNA editing efficiency scores significantly correlated with reporter editing, supporting the utility of this tool for prioritizing efficient pegRNAs in iPSCs **(Figure S6E).** However, reporter and endogenous editing were not significantly correlated due to confounding by chromatin accessibility, with pegRNAs targeting more accessible chromatin exhibiting a substantially higher ratio of endogenous to reporter editing compared to those targeting less accessible regions (**Figure S6F**). These findings are consistent with previous reports for both base and prime editing systems^51,56^.

We evaluated whether PE efficiency at low-accessibility chromatin in iPSCs could be enhanced by treatment with a histone deacetylase inhibitor (HDACi) given that HDACis have been previously shown to increase CRISPR/Cas9-mediated gene editing in various cell types^51,57,58^. iPSCs were transduced with the same pegRNA pools targeting either the highly accessible locus (region #1) or less accessible locus (region #2), with or without a 48 hour 1 mM valproic acid (VPA) treatment, and endogenous and reporter editing were measured over a four-week time course (**Figure 4B-C**). Treatment with VPA had no effect on editing efficiency at the highly accessible site (**Figure 4F**) or reporter (**Figure S6G**), but had increased editing efficiency of the less accessible site at week 3 **(Figure 4G).** These results demonstrate that treatment with VPA can increase prime editing efficiency at less accessible loci in iPSCs.

We next tested whether inhibition of mismatch repair (MMR) via expression of a dominant-negative MLH1 protein (hMLH1dn) could further improve editing efficiency across both loci, as has been previously demonstrated^59^. hMLH1dn was transduced into iPSCs two weeks after transduction of the pegRNA libraries and VPA treatment, and endogenous and reporter editing were measured after an additional two weeks, for a total of four weeks of editing **(Figure 4C)**. Stable expression of hMLH1dn significantly increased endogenous editing, although this effect did not reach statistical significance for the reporter (**Figure 4H**).

While VPA treatment and MMRi increased the correlation between endogenous and reporter editing **(Figure 4I)**, the ratio of endogenous to reporter editing remained strongly influenced by chromatin accessibility **(Figure S6H)**. To account for this, we fit a linear regression model incorporating reporter editing and ATAC-seq signal **(Figure S6H)**, and found that the resulting predicted editing efficiencies were highly correlated with observed endogenous editing (**Figure 4J)**. These results demonstrate that VPA and MMRi increase editing efficiency and, when combined with chromatin accessibility information, the reporter system can be used to better predict endogenous editing rates.

### PE HCR-FlowFISH screens identify common noncoding variants impacting expression of known SCZ genes

We compiled a list of variants to install and functionally characterize in the three WES-prioritized loci, including 1) common variants that map within CRISPRi-identified REs, 2) common variants in UTRs or splice sites of the WES gene, 3) SCZ FINEMAP variants, 4) 10 synthetic promoter indels within ±100 bp of the TSS as controls, and 5) 5 bp deletions centered on common SNPs from the previous categories to allow for detection of larger effect sizes. We conducted a SNP analysis to determine the SNP profile of our iPSC line, finding 97 prioritized variants where the cell line harbored either homozygous or heterozygous alternate variants relative to the hg38 reference genome (**Figure S7A**). Two representative homozygous ALT variants were validated by PCR amplification of the endogenous locus followed by Sanger sequencing (**Figure S7B**). We then used PRIDICT2.0 to generate pegRNAs tailored to the observed iPSC SNP profile.

We further narrowed the prime editing libraries by removing common variants that did not have either one pegRNA with at least moderate efficiency as predicted by PRIDICT2.0 or a prioritized functional impact in a disease-relevant genomic dataset^45,46,41^ (**Figure S7C, Table S11**). This resulted in a total of 2,386 pegRNAs (**Figure S7D**) installing 758 unique variants (**Figure S7E, Figure S8)** with up to 4 pegRNAs per common variant and up to 2 pegRNAs per synthetic indel. To control for the possibility of steric hindrance causing altered gene expression (independent of an edit), we included matched pegRNA controls installing the iPSC reference variant for each pegRNA, which we used for normalization in post-screen analysis **(Figure S7F, Methods)**. We ensured that no pegRNAs mutated the PAM nor gRNA seed regions given that this would alter the steric effect of a targeting pegRNA relative to its matched control pegRNA. Last, we included sets of 100-200 non-targeting pegRNAs as controls for all three libraries (**Tables S14-16**).

We transduced each library into iPSCs with VPA and hMLH1dn treatment given that these conditions improved editing efficiency in our previous validations (**Methods, Figure 4G, Figure S6H**). We performed HCR-FlowFISH screening three weeks post-transduction, sorting cells into bins based on low (0-20 percentile) or high (80-100 percentile) target gene expression (**Figure 5A**). Sequencing of the reporter in bulk unsorted cells revealed a median editing efficiency of 17-23% for each library **(Figure 5B),** which were correlated with PRIDICT2.0 scores (**Figure S7G**). The *STAG1* locus library displayed the lowest pegRNA reporter editing; and therefore, we additionally collected cells in the 20-40 and 60-80 percentiles to increase screen sensitivity^51^. We used BEAN^51^ to quantify variant effects on gene expression, normalizing for predicted endogenous editing efficiency and effect of each matched control pegRNA (**Methods**). As expected, 5 bp deletions surrounding the target genes’ TSSs led to larger decreases in gene expression relative to all other targeting pegRNAs and non-targeting controls (**Figure 5C**).

**Figure 5:**
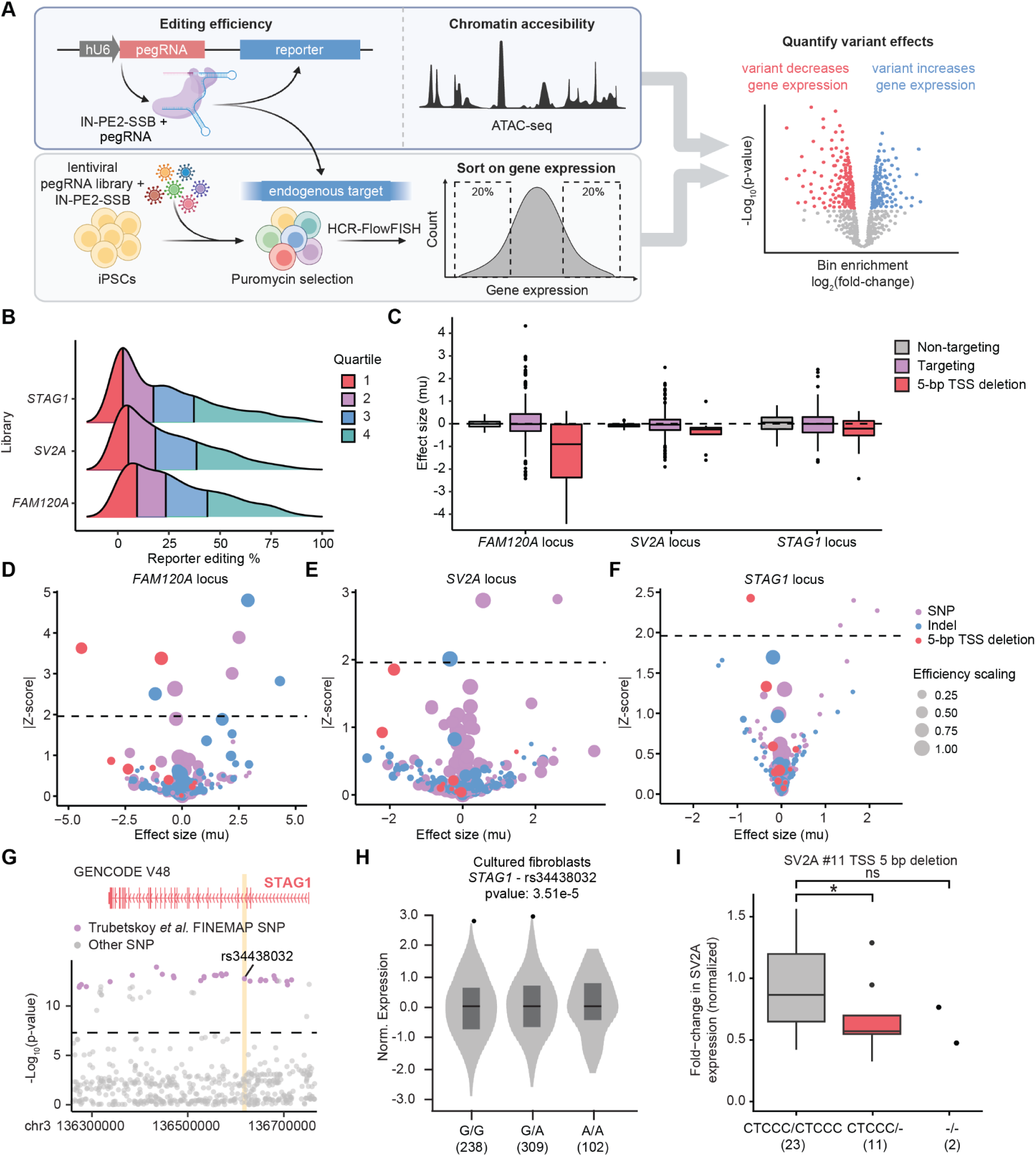
Activity-normalized prime editing (PE) screens identify variants impacting SCZ gene expression. A) Schematic of the PE screening strategy. IN-PE2-SSB editor and pegRNA libraries are transduced into iPSCs, which are later treated with VPA and transduced with hMLH1dn. Cells are sorted into bins (bottom 20% and top 20%) based on target gene expression via HCR-FlowFISH. Variant effects are calculated using BEAN^51^, with reporter editing and chromatin accessibility used to scale variant effect sizes. B) Distribution of the percent reporter editing for pegRNAs in the three PE screens. Vertical lines depict quantiles. C) BEAN effect sizes for non-targeting, targeting, and 5 bp TSS deletion edits in the three PE screens. Volcano plots depicting BEAN effect sizes and Z-scores for variants in the (D) *FAM120A* locus, (E) *SV2A* locus, or (F) *STAG1* locus. G) Locus view displaying SCZ variants in the region surrounding the *STAG1* gene body. SCZ FINEMAP SNP rs34438032:G>A is highlighted in yellow in the third intron of *STAG1*. H) Violin plot of normalized *STAG1* expression across rs34438032 genotypes (GG, GA, AA) in cultured fibroblasts from GTEx. I) Normalized *SV2A* expression in iPSC 5 bp SV2A TSS deletion #11 edited clones across three genotypes as measured by RT-qPCR. Number of clones in parentheses. Two-tailed t test. (∗p < 0.05, ∗∗p ≤ 0.01, ∗∗∗p ≤ 0.001).

We quantified variant effects on expression of the three SCZ genes, with significant variant hits including three 5 bp TSS deletions, eight SNPs, three 5 bp synthetic deletions centered on SNPs, and one insertion (|Z-score| >= 1.98, **Figure 5D-F, Tables S17-19**). Of the 11 SNPs prioritized through direct installation or 5 bp deletion, only five of these variants were imputed in the SCZ GWAS study given MAFs < 1% in 1000 Genomes EUR for the other variant hits^60^. The top *STAG1* SNP hit, rs34438032:G>A is a SCZ FINEMAP SNP^5^, making this a highly prioritized variant (**Figure 5G)**. Interestingly, this variant does not map in open-chromatin in iPSCs nor iNeurons, so this region was not identified as a RE in our CRISPRi screens. This variant is associated with an increase in *STAG1* expression in cultured fibroblasts in GTEx (**Figure 5H**), consistent with the increase in *STAG1* expression observed in the PE screen. In the *FAM120A* locus, a FINEMAP SNP in the promoter of *FAM120A*, rs7853605:A>G (**Figure 3C**), significantly increased expression of this gene, in line with GTEx eQTL data from esophagus - mucosa (**Figure 3D**). Another common EUR variant hit, rs112851681:A>G, was located within the *SV2A* promoter (**Figure S8**), but exhibited low LD with the index SNP, and is therefore unlikely to drive the association signal detected in the SCZ GWAS. Nevertheless, this SNP may represent a secondary association signal that remains underpowered in existing studies. The six SNPs that were rarer in European populations may contribute more substantially to SCZ risk in other populations where they are more common.

### Endogenous validation of functional noncoding variants

To validate variants prioritized through the pooled PE screens, we transiently transfected three pegRNA constructs into iPSCs, isolated a total of 115 clones, and performed genotyping and RT-qPCR. We observed a significant decrease in *SV2A* expression for heterozygous clones harboring the top *SV2A* 5 bp promoter deletion, pegSV2A #11 (**Figure 5I**). We observed a trending but non-significant increase in *SV2A* expression and decrease in *FAM120A* expression for heterozygous rs112851681:A>G and rs117810130:C>T clones, respectively (**Figure S7H**), in line with our pooled PE screen results (**Figure 5E**). We did not observe significant effects for homozygous edited clones due to the limited sample size for these genotypes (**Figure 5G, Figure S7H**). We note that this approach lacks statistical power, as clonal isolation of edited cells is low throughput, variant effects in their endogenous context are likely subtle relative to CRISPRi of the RE, and clone-to-clone variation can result in large variability of gene expression.

### GFP reporter for validation of variant effects on enhancer activity in iPSCs and iNeurons

To validate variant effects on enhancer activity using a more sensitive and scalable approach, we individually cloned 200 bp sequences including either reference (REF) or alternate (ALT) alleles into a lentiviral GFP reporter construct containing a minimal promoter (**Figure 6A**). We transduced reporter constructs into iPSCs at an MOI of 1.5 and measured GFP expression for eight variant hits and two variant non-hits (**Figure 6A**). We found that the *SV2A* 5 bp deletion #11 **(Figure 6B)** and all tested SNP PE screen hits had significant impacts on enhancer activity in the same direction as the pooled PE screen **(Figure 6C)**. Also in line with the PE screens, we did not observe significant impacts on enhancer activity for the two non-hits (**Figure S9A**). However, two 5 bp synthetic deletion hits, as well as their corresponding non-hit SNPs, did not cause significant impacts on enhancer activity in the reporter context (**Figure S9A**). Interestingly, these non-validating variants were located in elements with decreased baseline enhancer activity compared to those showing significant effects (**Figure S9B**). This could suggest that these variants require additional sequence context to impact transcriptional activity, either because their effects depend on a larger portion of the RE than the ∼200 bp sequence tested in the reporter assay or through interactions with other REs.

**Figure 6:**
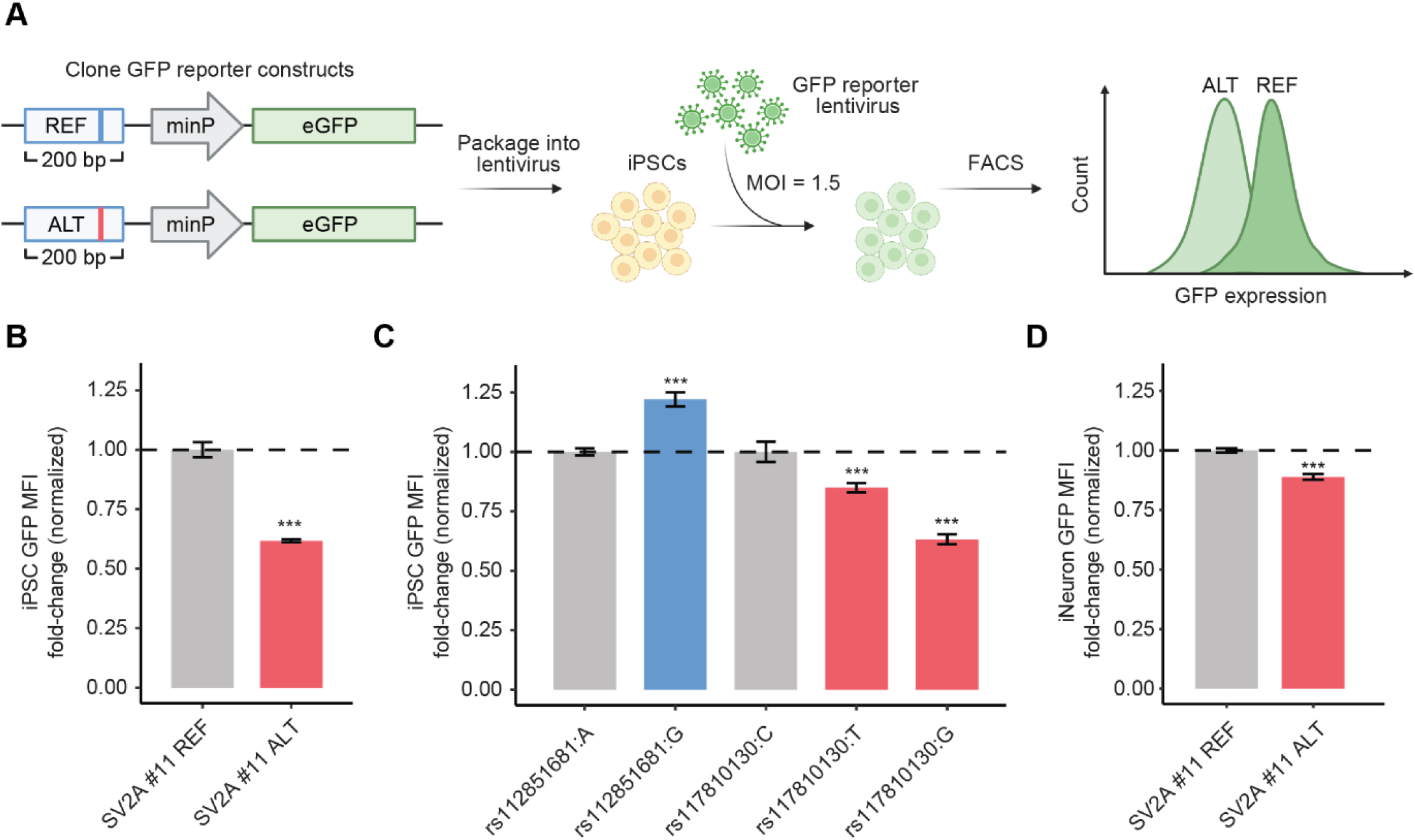
A GFP reporter quantifies variant effects on enhancer activity in iPSCs and iNeurons. A) Schematic of the GFP reporter assay. 200 bp sequences containing reference (REF) or alternative (ALT) variants are cloned in front of a minimal promoter (minP) driving GFP expression. These constructs are packaged into lentivirus and transduced into cells at an MOI of 1.5. The relative levels of GFP are then assayed using FACS. B) Normalized fold-change in GFP MFI for the SV2A #11 5 bp deletion relative to the reference sequence in iPSCs. n=6. Two-tailed t test. C) Normalized fold-change in GFP MFI for three ALT SNPs relative to reference SNPs in iPSCs. n=4-6. Two-tailed t test. D) Normalized fold-change in GFP MFI for the SV2A #11 5 bp deletion relative to the reference sequence in iNeurons. n=3. Two-tailed t test. (∗p < 0.05, ∗∗p ≤ 0.01, ∗∗∗p ≤ 0.001).

We next tested whether the GFP reporter system could be used to identify variants impacting enhancer activity in iNeurons. We found that the *SV2A* 5 bp deletion #11 resulted in decreased enhancer activity in D7 iNeurons (**Figure 6D**), however, this effect was of a smaller magnitude in iNeurons relative to iPSCs (**Figure 6B**). This may be due in part to transgene silencing during the iNeuron differentiation^61^. We observed significantly reduced enhancer activity of the promoter sequence in iNeurons compared to iPSCs, with a large portion of the transduced population overlapping the spectrum of non-transduced cells (**Figure S9B-D**), even though *SV2A* is more highly expressed in D7 iNeurons (**Figure S1F**). This indicates that assessing variant effects using a reporter system is possible in iNeurons, however, for some targets this approach could have reduced sensitivity relative to iPSCs.

### rs112851681:G increases transcriptional activity in iNeurons

We next investigated how variants identified in iPSC pooled PE screens impact expression in iNeurons. rs112851681:G is a SNP in the promoter of *SV2A* (**Figure S8C**) that increased expression of *SV2A* in the pooled PE screen (**Figure 5E**) and increased regulatory element activity in iPSCs **(Figure 6C**). We also show that rs112851681:G significantly increases enhancer activity in iNeurons (**Figure 7A**). In line with this, rs112851681:G results in an increase in sequence similarity to multiple motifs in the SP-family (**Figure 7B**), all of which have detectable expression in iPSCs and iNeurons (**Table S2**). This family of TFs includes the TF encoded by *SP4*, a known SCZ risk gene that is predominantly expressed in neurons^62,63^ and that has been shown to activate gene expression by binding to promoters^64^. Concordantly, rs112851681:G is associated with a significant increase in expression of *SV2A* in GTEx Cerebellar Hemisphere tissue (**Figure 7C)**, a tissue containing a large population of excitatory glutamatergic neurons^65,66^. We performed RT-qPCR of *SV2A* on 33 clonal cell lines differentiated into D7 iNeurons and containing distinct rs112851681 genotypes and observed a ∼25% increase in SV2A expression for heterozygous clones (p=0.1, **Figure 7D**). Taken together, these results suggest that rs112851681:G can influence *SV2A* expression through increased transcriptional activity, likely mediated through increased binding of one or more TFs.

**Figure 7:**
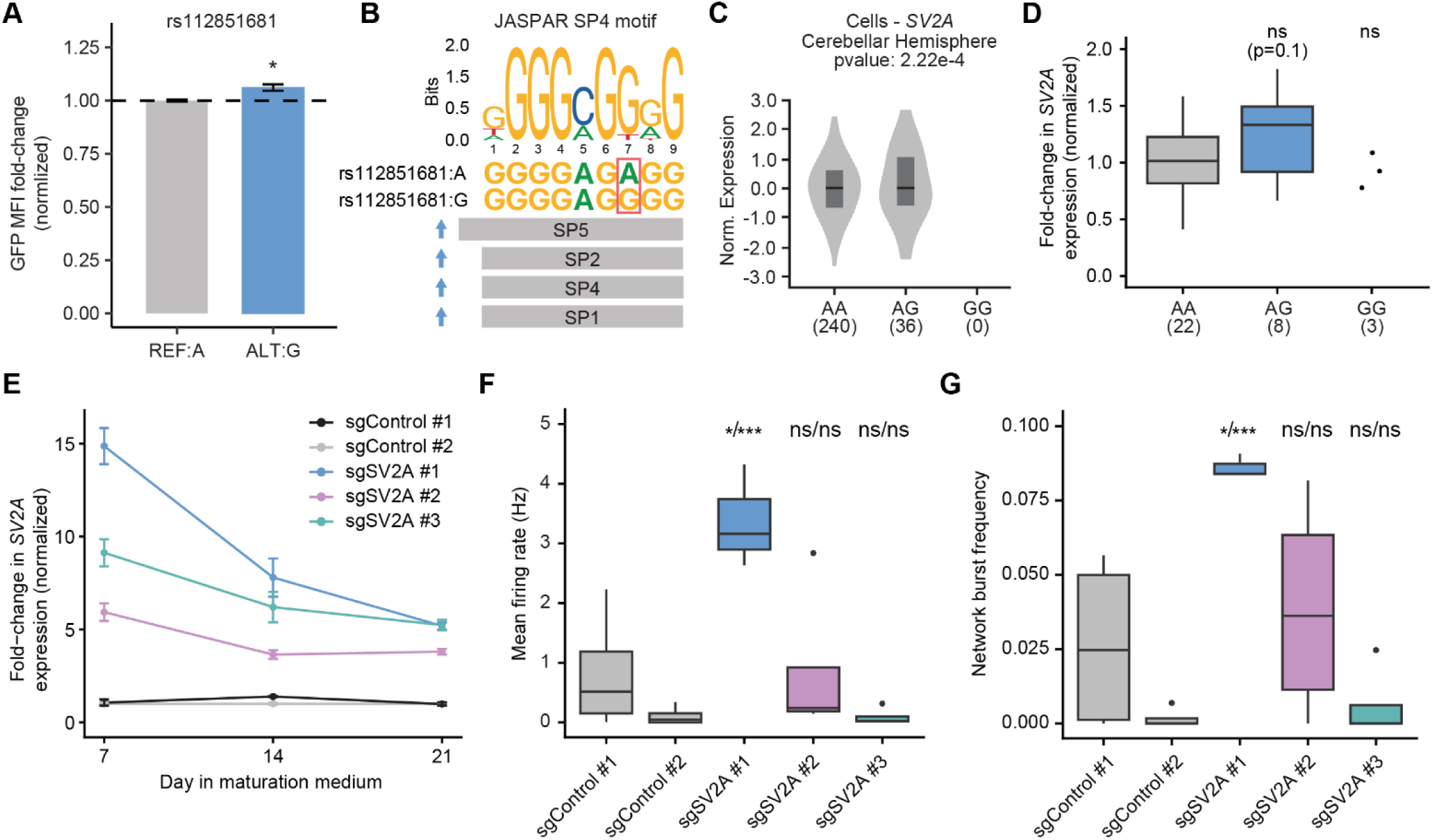
rs112851681:G increases transcriptional activity in iNeurons. A) Normalized fold-change in GFP MFI for rs112851681:G relative to rs112851681:A in iNeurons. n=3. Two-tailed t test. B) SP family TF position weight matrices (PWMs) impacted by rs112851681. C) Violin plot of normalized *SV2A* expression across rs112851681 genotypes (AA, AG, GG) in cerebellar hemisphere from GTEx. D) Normalized *SV2A* expression in D7 iNeuron rs112851681 edited clones across three genotypes (AA, AG, GG) as measured by RT-qPCR. Two-tailed t test. E) Normalized *SV2A* expression of DHFR-dCas9-VPH iNeurons transduced with either non-targeting control or *SV2A* gRNAs over a 3 week time course as measured by RT-qPCR. n=3. F) Mean firing rate of D21 DHFR-dCas9-VPH iNeurons transduced with either non-targeting control or *SV2A* gRNAs. n=3-4. Two-tailed t test. G) Network burst rate of D21 DHFR-dCas9-VPH iNeurons transduced with eithernon-targeting control or *SV2A* gRNAs. Two-tailed t test. (∗p < 0.05, ∗∗p ≤ 0.01, ∗∗∗p ≤ 0.001).

### SV2A upregulation increases neuronal network activity

While disruptive coding variants in *SV2A* are associated with increased risk of SCZ^5^, increased *SV2A* expression has yet to be linked to this disorder. Previous studies have shown that both *SV2A* knockout and *SV2A* exogenous overexpression produce parallel phenotypes reflecting reduced synaptic release probability^67,68^. To determine if increasing *SV2A* levels alters network activity in iNeurons, we performed a multielectrode array (MEA) assay with CRISPRa of *SV2A*. We tested three CRISPRa gRNAs (sgSV2A #1-3) targeting the promoter of *SV2A*, observing significant upregulation of *SV2A* for all three gRNAs throughout neuronal differentiation relative to two non-targeting gRNAs (sgControl #1-2) (**Figure 7E**). Interestingly, *SV2A* upregulation was relatively stable for iNeurons expressing sgSV2A #2 and #3, while iNeurons expressing sgSV2A #1 had the largest upregulation at day 7, and then reduced their upregulation over time (**Figure 7E**). This indicates possible feedback mechanisms to reverse strong *SV2A* upregulation. Nevertheless, iNeurons expressing sgSV2A #1, which had the most robust *SV2A* activation overall, displayed a significantly increased mean firing rate (**Figure 7F**), burst rate (**Figure S10A**), and network burst rate (**Figure 7G**) relative to iNeurons expressing non-targeting control gRNAs. These data demonstrate that increased *SV2A* expression can alter network activity in iNeurons and further support that aberrant *SV2A* expression outside of typical levels can impact synaptic transmission.

## Discussion

In this study, we combined WES^6^, GWAS^5^, and CRISPR-based functional genomics in iPSCs and iNeurons to dissect the function of REs and variants implicated in SCZ risk. While statistical genetics has uncovered both rare and common variants that contribute to disease, moving from association signals to mechanistic insight has remained a major challenge. By applying high-throughput CRISPRi and PE screens at loci overlapping WES and GWAS signals, we systematically mapped REs and variants impacting expression of three WES-prioritized SCZ genes. This framework not only strengthens causal inferences at these loci, but also provides a pipeline for investigating the genetic architecture of other SCZ and disease-associated regions.

Our RNA-seq and ATAC-seq datasets across the iPSC-to-neuron differentiation time course provide a valuable resource for nominating loci and genes relevant to neurodevelopmental disorders that can be investigated using these cellular models. The ATAC-seq dataset identifies pREs present in iPSCs and iNeurons within disease-associated loci. The RNA-seq time course defines the expression patterns of genes in iPSCs and iNeurons, pinpointing when disease relevant genes are most highly expressed during the iPSC-to-neuron differentiation. Together, these datasets aid in establishing the regulatory landscape of neuronal models to increase their utility in deciphering the genetic basis of complex neurodevelopmental and neuropsychiatric disorders.

We established a pipeline for screening disease-relevant pREs in iPSCs and iNeurons using CRISPRi HCR-FlowFISH. This pipeline is scalable and can be implemented across diverse loci, gene targets, and neurodevelopmental diseases to map REs influencing gene expression. We assayed 333 pREs across three WES-prioritized loci, identifying 78 REs that significantly modulate the expression of SCZ genes *FAM120A*, *STAG1*, and *SV2A*, and prioritizing over a thousand common variants from diverse population studies for further investigation. The CRISPRi-identified REs included both proximal and distal regulators, with some located hundreds of kb away from the promoters of WES genes. Overlapping CRISPRi-identified REs with eHi-C datasets supports putative *cis*- and *trans*-mechanisms of effect, but further investigation is needed to validate these hypothesized mechanisms and determine the factors modulating their effects. These results demonstrate the complexity of gene regulation in SCZ-associated loci, where REs can simultaneously influence expression of multiple genes and/or noncoding RNAs.

Overlapping CRISPRi-identified regulatory elements with functional genomic datasets, including xQTLs, MPRA results, and FINEMAP-prioritized SNPs generated additional support for SCZ-associated variants mapping in REs (**Table S11**). This approach connected noncoding variation to gene-expression changes, informed hypotheses about underlying mechanisms, and prioritized regions that may contain variants that impact regulatory element function. However, our ability to use genetic datasets to interpret the effect of common noncoding indels, variants in underrepresented populations, and variants in high LD remains limited.

To address the current limitations in noncoding variant interpretation, we optimized and implemented PE screens in iPSCs to investigate the impact of disease-associated variants. We found 15 variants that significantly impact the expression of a known SCZ gene, including eight noncoding SNPs from multi-ancestry cohorts. We validated variant effects using RT-qPCR and a GFP reporter system in iPSCs and iNeurons. We found that rs112851681:G is a common noncoding SNP in the promoter of *SV2A* that increases transcriptional activity in iPSCs and iNeurons. We provided orthogonal evidence for the role of this variant in modulating *SV2A* expression by integrating eQTL data and predicting the effect of alternate alleles on TF binding motifs, including the predicted creation of an SP-family binding motif. While robust CRISPRa of *SV2A* resulted in altered firing of neuronal networks, the impact of variants with subtle influences on *SV2A* expression will be important to investigate.

Our CRISPR screens demonstrate that REs and noncoding variants in SCZ-associated GWAS loci influence gene expression of *FAM120A*, *STAG1*, and *SV2A in vitro*. Generating whole genome sequencing data for SCZ patients from diverse ancestries would provide additional lines of evidence to support the role of these variants in disease risk, particularly for indels and variants not common in European populations. We also note that REs and variants that impact expression *in vivo* but that do not have effects in iPSCs and iNeurons may be false negatives in our screens. Nevertheless, we show that this framework identifies noncoding mechanisms that lead to differential expression of disease genes in disease-relevant cell models.

This work provides a deep interrogation of the role of the noncoding genome in regulation of three known SCZ genes, providing a template for the study of other complex polygenic trait and disease loci using high-throughput CRISPR screening methodologies. As GWAS and WES studies continue to identify risk loci across neuropsychiatric disorders, neurodevelopmental disorders, and beyond, this framework offers a scalable route from statistical association toward mechanistic understanding of disease-associated variants.

## Acknowledgments

We thank the Duke cell culture facility for excellent assistance. We also thank the teams at High-throughput Applied Research Data Analysis Cluster (HARDAC) and Duke Computing Cluster (DCC) for computing resources. We thank Luca Pinello and Ann Ciricione for their expertise and support in the BEAN analysis. We thank Ellie Guilfoyle and Carson Key for their input on the GFP reporter assays. Schematics were created with BioRender.com. The Genotype-Tissue Expression (GTEx) Project was supported by the Common Fund of the Office of the Director of the National Institutes of Health, and by NCI, NHGRI, NHLBI, NIDA, NIMH, and NINDS. The GTEx data used for the analyses described in this manuscript were obtained from the GTEx Portal on 04/29/24. The work is funded by UM1-HG012053, RM1-HG011123 and R01-MH125236 and Open Philanthropy Project. M.C.H. was supported by the NIH-F31 (F31MH138142).

## Author contributions

Conceptualization, methodology, writing – original draft, and writing – reviewing and editing, M.C.H, A.C.N, J.W.R, B.L., R.I.S, M.I.L., A.S.A., P.F.S., C.A.G., G.E.C.; investigation and validation, M.C.H, A.C.N, J.W.R, B.L., S.F.D, A.S., A.A.C, K.T.H., A.B., P.F.S.; software, formal analysis and visualization, M.C.H, A.C.N, J.W.R, B.L., A.A.C, A.B., R.R., M.I.L., P.F.S.; funding acquisition and supervision, C.A.G., G.E.C., P.F.S., M.C.H, J.W.R.

## Inclusion and Diversity

We support inclusive, diverse, and equitable conduct of research.

Supplemental_Tables

**Supplemental Table 1:** Summary of loci with SCZ-GWAS/WES overlap, Related to Figure 1

**Supplemental Table 2:** RNA-seq, Related to Figure 1

**Supplemental Table 3:** ATAC-seq, Related to Figure 1

**Supplemental Table 4:** *FAM120A* locus gRNA library information, Related to Figure 2

**Supplemental Table 5:** *STAG1* locus gRNA library information, Related to Figure 2

**Supplemental Table 6:** *SV2A* locus gRNA library information, Related to Figure 2

**Supplemental Table 7:** *FAM120A* locus iPSC CRISPRi screen summary, Related to Figure 2

**Supplemental Table 8:** *STAG1* locus iPSC CRISPRi screen summary, Related to Figure 2

**Supplemental Table 9:** *SV2A* locus iPSC CRISPRi screen summary, Related to Figure 2

**Supplemental Table 10:** *SV2A* locus iNeuron CRISPRi screen summary, Related to Figure 2

**Supplemental Table 11:** Prioritized SCZ GWAS variants, Related to Figure 3

**Supplemental Table 12:** Region #1 pegRNA library information, Related to Figure 4

**Supplemental Table 13:** Region #2 pegRNA library information, Related to Figure 4

**Supplemental Table 14:** *FAM120A* locus pegRNA library information, Related to Figure 5

**Supplemental Table 15:** *STAG1* locus pegRNA library information, Related to Figure 5

**Supplemental Table 16:** *SV2A* locus pegRNA library information, Related to Figure 5

**Supplemental Table 17:** *FAM120A* locus prime editing screen summary, Related to Figure 5

**Supplemental Table 18:** *SV2A* locus prime editing screen summary, Related to Figure 5

**Supplemental Table 19:** *STAG1* locus prime editing screen summary, Related to Figure 5

## Resource Availability

### Lead contact

Further information and requests for resources and reagents should be directed to and will be fulfilled by the lead contacts, Charles A. Gersbach, Gregory E. Crawford, and Patrick Sullivan.

### Materials availability

Plasmids generated in this study have been deposited to Addgene and will be publicly available as of the date of publication.

### Data and code availability

IGVF data portal accession numbers will be provided before the date of publication.

## Experimental Model and Subject Details

### Cell lines

The i^3^N WTC-11 iPSC lines containing an inducible mNgn2 knock-in cassette in the AAVS1 safe-harbor locus and expressing dCas9^KRAB^ or no CRISPR machinery were gifts from Dr. Yin Shen’s lab at UCSF^12,69^. The i^3^N WTC-11 iPSC line containing an inducible mNgn2 knock-in cassette in the AAVS1 safe-harbor locus and expressing DHFR-dCas9-VPH was a gift from Dr. Martin Kampmann’s lab at UCSF^70^.

## Method Details

### Cell lines and culture conditions

All cells were grown at 37°C + 5% CO2. HEK293T/17 cells were cultured in DMEM + 10% FBS. WTC-11 cells were cultured on matrigel coated plates and were maintained in complete mTeSR Plus medium (StemCell Tech, 100–0276).

### ATAC-seq Cell Processing and Library Preparation

Cultured cells at designated time points at iPSC stage (Dm4 - 2 biological replicates) and throughout differentiation (D0/D3/D7/D14/D21/D28 - 3, 2, 3, 3, 1, and 3 biological replicates each) were first washed with DPBS, harvested by Accutase treatment, for Dm4 and D, or by scraping or Accutase/papain treatment (for details of this method, see **Methods** subsection “CRISPRi HCR-FlowFISH screen tissue culture”), for the rest of the time points, into DPBS, and pelleted by centrifugation at 200–300 x g for 5 minutes at room temperature. Cell pellets were then resuspended in cryopreservation media (90% KnockOut Serum Replacement (Gibco, 10828010) + 10% dimethyl sulfoxide (Sigma-Aldrich, D8418)) and stored at −80°C.

Approximately 100,000-500,000 viable frozen cells were used for ATAC-seq library preparation according to the Omni-ATAC protocol (*93*) with modifications. Frozen vials were thawed at 37°C. Cells were washed with warm growth media then washed with 1X PBS. Cell pellets were suspended in 1mL of ATAC-seq RSB containing 0.1% Tween and transferred to a dounce homogenizer. Cells were lysed with 10 stokes (tight pestle) and centrifuged for 10 min at 500 rcf. Nuclei pellets were suspended in 27μl 2X TD buffer and 2ul were used for counting using hemocytometer. 50,000 nuclei were added to the transposition reaction mix (2.5 μl transposase, 16.5 μl PBS, 0.5 μl 1% digitonin, 0.5 μl 10% Tween-20, 5 μl water, and 2X TD buffer to 50ul) and incubated at 37°C for 30 min in a thermomixer with shaking at 1,000 rpm. Reactions were cleaned up with Zymo DNA Clean and Concentrator-5 kit. ATAC-seq libraries were double indexed with Nextera PCR Primers and amplified with 9 to 12 cycles of PCR. Amplified DNA fragments were purified with 1:1 ratio of Agencourt AMPure XP (Beckman Coulter). Libraries were quantified by Qubit and size distribution was inspected by Tapestation (Agilent High Sensitivity DNA, Agilent Technologies).

After ATAC-seq library generation, library yield and size distributions were assessed using a Qubit dsDNA BR Assay and an Agilent TapeStation with D5000 HS screen tapes, respectively. Individual libraries were normalized to 4 nM and pooled at equimolar ratios. The final library pool was further diluted to 500 pM using Resuspension Buffer (RSB) with Tween, and paired-end sequencing was performed on an Illumina NextSeq instrument. An average of 55 million paired-end reads per library were obtained.

Additional D3 and D14 samples (1 biological replicate each) were dissociated and prepared for ATAC-seq as follows. 25 μL of DNase I (Sigma-Aldrich, DN25, 400 U/ml, 400 Kunitz unit/mg protein) and 5 μL of 1M MgCl2 were added to each 1 mL of activated papain and the tube was inverted twice to thoroughly mix reagents. Media was removed from human 2-week neurons in 12-well plates and cells were carefully washed once with 1x HBSS (200 μL/well). Then, 200 μL of activated papain was added per well and cells were incubated at 37°C for 30 mins. After dissociation, cells were quenched with 600 μL of DMEM/F12 medium (Gibco, 11330–0322) containing 10% Fetal Bovine Serum (FBS) (Sigma-Aldrich, 9628642). All neurons were then collected from each well in a 50 mL tube and triturated 10–15 times with a 1-mL pipette to make them into single cells.

Cell were prepared as previously described using the Nextera DNA Library Prep Kit (Illumina #FC-121–1030). Briefly, fixed cells were washed once with ice cold PBS containing 1x protease inhibitor before being resuspended in ice cold nuclei extraction buffer (10 mM Tris-HCl pH 7.5, 10 mM NaCl, 3 mM MgCl2, 0.1% Igepal CA630, and 1x protease inhibitor) for 5 minutes. 50,000 cells were aliquoted, exchanged into 50 μL 1x Buffer TD, and incubated with 2.5 μL TDE1 enzyme for 45 minutes at 37°C with shaking. Following transposition, 150 μL reverse crosslinking solution (50 μL 1 M Tris pH 8.0, 100 μL 10% SDS, 2 μL 0.5 M EDTA, 10 μL 5 M NaCl, 800 μL water, and 2.5 μL 20 mg/mL Proteinase K) was added to each tube and incubated at 65°C overnight. DNA was column purified, PCR amplified, and size-selected for fragments between 300 and 1000 bp. Libraries were sent for paired-end sequencing on the NovaSeq 6000 instrument (150 bp paired-end reads).

### ATAC-seq Data Processing

Raw ATAC-seq reads for 21 biological samples (19 generated in this study, plus 2 published (ENCLB066COW/ENCLB316BUG - D28; ENCLB745RLS/ENCLB868CJL - Dm4)) were mapped to human reference GRCh38 and processed with the ENCODE ATAC-seq pipeline (https://github.com/ENCODE-DCC/atac-seq-pipeline) (PMID: 37066421) that is adapted to output peak calls with FDR <0.01 instead of the default IDR method. A reproducible union peak set was called across all samples with MSPC (PMID: 25957351), requiring a peak to be called in at least two samples to be kept for downstream analysis. This resulted in a set of 314,870 peaks. Finally, featureCounts (PMID: 24227677) was used to count the numbers of reads overlapping the peaks for each sample.

### RNA-seq Cell Processing and Library Preparation

Cultured cells at designated time points were harvested and cryoperserved as described in **Methods** subsection “ATAC-seq Cell Processing and Library Preparation”. To prepare RNA, cryopreserved cell aliquots were thawed in a 37°C water bath. To maximize cell recovery and minimize osmotic shock, pre-warmed DMEM/F12 media was added to the cell suspension slowly and dropwise while swirling. The diluted cell suspensions were then pelleted by centrifugation at 300 x g for 5 minutes. The supernatant was carefully decanted, and the resulting cell pellets were immediately lysed in Buffer RLT (using 350 µL per standard pellet, and up to 700 µL for denser samples). Total RNA was extracted using the RNeasy Kit following the manufacturer’s standard protocol (RNeasy Mini Kit, Qiagen 74104), without on-column DNA digestion. Purified RNA was eluted in 30 µL of RNase-free water. Following extraction, total RNA yields and sample purity were quantified using a Qubit RNA BR assay.

RNA samples were then shipped to Genewiz for RNA-seq library prep (Library preparation, Illumina, RNA with rRNA depletion) and sequencing (Illumina, 2×150bp, ∼350M PE reads) services, with targeted sequencing depth at 30 million paired-end reads per sample.

### RNA-seq Data Processing

Raw RNA-seq reads for 18 biological samples (all generated in this study) were mapped to human reference GRCh38 (gene annotation Gencode v41) and processed with the ENCODE RNA-seq pipeline (https://github.com/ENCODE-DCC/rna-seq-pipeline) (PMID: 37066421). Transcript Per Million (TPM) values were taken from the gene-level RSEM output files and combined across samples from the same time points.

### CRISPRi library design and cloning

For each SCZ GWAS locus library, up to 20 gRNAs were designed targeting the union set of ATAC-peaks mapping within each GWAS locus (loci extended by 10%) from the iPSC to iNeuron differentiation timecourse. gRNAs were selected using GuideScan2 with a specificity threshold of 0.1 to minimize off-target effects. gRNAs harboring a TTTT sequence were removed from the gRNA library. 200 non-targeting control gRNAs were included per locus. All oligonucleotide libraries (**Tables S4-6**) were ordered in the following sequence format:

FAM120A: 5’ TGATCAATCCGCGCCATGAC ATATATCTTGTGGAAAGGACGAAACACCG [20-bp protospacer] GTTTAAGAGCTATGCTGGAAACAGCATAG 3’

STAG1: 5’ TTGGATGCAGGTCGAAAGGC ATATATCTTGTGGAAAGGACGAAACACCG [20-bp protospacer] GTTTAAGAGCTATGCTGGAAACAGCATAG 3’

SV2A: 5’ CCTTGTCAACAGACCATGCC ATATATCTTGTGGAAAGGACGAAACACCG [20-bp protospacer] GTTTAAGAGCTATGCTGGAAACAGCATAG 3’

Libraries were amplified by PCR using Q5® High-Fidelity 2X Master Mix (NEB) using the following primers and annealing temperatures:

PCR1:

FAM120A_sublib_fw: 5’ TGATCAATCCGCGCCATGAC 3’ Ta = 69 °C

STAG1_sublib_fw: 5’ TTGGATGCAGGTCGAAAGGC 3’ Ta = 69 °C

SV2A_sublib_fw: 5’ CCTTGTCAACAGACCATGCC 3’ Ta = 67 °C

gRNA_60bp_rv 5’ GTTGATAACGGACTAGCCTTATTTAAACTTGCTATGCTGTTTCCAGCATAGCTCTTAAAC 3’

PCR2:

gRNA_60bp_fw 5’ TAACTTGAAAGTATTTCGATTTCTTGGCTTTATATATCTTGTGGAAAGGACGAAACACCG 3’

gRNA_60bp_rv 5’ GTTGATAACGGACTAGCCTTATTTAAACTTGCTATGCTGTTTCCAGCATAGCTCTTAAAC 3’

Ta = 72 °C

gRNA libraries were cloned into a lentiviral gRNA expression vector with BFP-P2A-PuroR (Addgene plasmid #230935) through BsmBI vector digest and NEBuilder HiFi DNA assembly, ensuring >100-fold representation of each gRNA.

### CRISPRi HCR-FlowFISH screen tissue culture

For WTC-11 dCas9^KRAB^ cell screens, cells were seeded at a density of 4.34×10^4^ cells/cm^2^ on 10-cm plates in four biological replicates in the presence of 10 µM Y-27632. The cells were transduced with lentivirus at a multiplicity of infection (MOI) of 0.3 as determined by titration. Two days post-transduction, cells were treated with 0.5 µg/mL puromycin and were selected for 8 days. After selection, cells were lifted off the plate using accutase and used as input for genomic DNA (gDNA) isolation or the HCR-flowFISH protocol detailed below.

For iNeuron dCas9^KRAB^ screens, WTC-11 dCas9^KRAB^ cells were seeded at a density of 1.15×10^5^ cells/cm^2^ on 15-cm plates in four biological replicates in the presence of 10 µM Y-27632 on day −4. The following day, media was changed to pre-differentiation media (PDM; Knockout DMEM/F-12 (1X) supplemented with N-2 (1X), NEAA (1X), BDNF (10ng/mL), NT-3 (10ng/mL), laminin (1 µg/mL), and doxycycline (2 µg/mL)), which was refreshed each day for three total days. Day 0 neurons were seeded at a density of 1.6×10^5^ cells/cm^2^ on three 15-cm plates per replicate in four biological replicates in maturation medium (Neurobasal-A (0.5X), DMEM/F12-HEPES (0.5X), NEAA (1X), B-27 (0.5X), N-2 (0.5X), GlutaMAX (0.5X), BDNF (10 ng/mL), NT-3 (10 ng/mL), laminin (1 µg/mL), and doxycycline (2 µg/mL)). The cells were transduced with lentivirus at a multiplicity of infection (MOI) of 0.6 as determined by titration. Seven days post-transduction, cells were lifted off the plate using an accutase/papain solution at 20 units of papain per mL, quenched in papain quench solution (DMEM F/12 + 1X GlutaMAX + 10 µM Y-27632 + 66.67 units Deoxyribonuclease I (DNase)), and used as input for gDNA isolation or the HCR-flowFISH protocol detailed below.

### HCR-FlowFISH

Fixation and Permeabilization: Cells were pelleted at 600 x g and resuspended in fixation/permeabilization buffer at a ratio of 1 mL per 5 million cells. Samples were incubated at room temperature for 60 minutes with gentle rotation. After fixation, cells were pelleted at 800 x g for 5 minutes, aspirated, and washed with PBS containing 0.1% Tween-20 (PBST). This wash step was repeated for a total of four washes. Following the final wash, cells were pelleted again at 800 x g and resuspended in the remaining PBST volume. Cells were transferred to ice and resuspended in pre-chilled 70% ethanol for 10 minutes. Cells were then pelleted at 1000 x g for 5 minutes and processed immediately for hybridization.

Hybridization and Signal Amplification: Hybridization buffer was thawed and pre-warmed to 37°C. Cells were washed twice with PBST and centrifuged at 800 x g for 5 minutes each time. The cell pellet was resuspended in hybridization buffer and pre-hybridized at 37°C for 30 minutes on a rotating mixer. During pre-hybridization, probe mixtures were prepared using probes targeting TBP and the target gene (Molecular Instruments) at the concentrations described below. The probe solution was added to each sample to achieve a 4 nM solution of probe. Samples were incubated overnight at 37°C on a rotating mixer.

Probe Washing and Hairpin Amplification: The next morning, probe wash buffer was thawed and pre-warmed to 37°C. Probe wash buffer was added to each tube, followed by centrifugation at 800 x g for 15 minutes. The supernatant was aspirated and cells were then resuspended in additional probe wash buffer. This wash step was repeated three more times, for a total of four washes. Clumps were broken up with a pipette if necessary.

After washing, cells were resuspended in saline-sodium citrate with 0.1% Tween-20 (5X SSCT) and incubated at room temperature for 5 minutes. Meanwhile, hairpins and amplification buffer were thawed at room temperature, protected from light. A heat block was preheated to 95°C. Hairpins (h1 and h2 for each fluorophore) were snap-cooled and added to the amplification buffer to reach a final concentration of 60 nM per hairpin. Cells were pelleted and resuspended in pre-warmed amplification buffer. After a 30-minute pre-amplification incubation at room temperature with rotation, the hairpin solution was added to each cell sample. Amplification was carried out overnight in the dark at room temperature with gentle rotation.

Post-Amplification Washing and Flow Cytometry: Following amplification, 5X SSCT was added to each sample. Cells were centrifuged at 800 x g for 5 minutes, and the supernatant was aspirated. The pellet was gently washed with 5X SSCT, followed by another spin at 800 x g for 5 minutes. This wash step was repeated five additional times (six total washes). Finally, cells were resuspended in FACS suspension media and passed through a 35 µm cell strainer three times to minimize clumps. Prepared samples were analyzed by flow cytometry.

### CRISPRi HCR-FlowFISH screen sorting and NGS

Cells were sorted into two bins (bottom 15% and top 15%) using a Sony Sorter flow cytometer. After sorting, gDNA was isolated from cells and was used to amplify the U6-3’ to gRNA hairpin region with different in-line barcodes. PCR2 was performed to add full-length Illumina sequencing adapters using internally ordered primers with equivalent sequences to NEBNext Index Primer Sets 1 and 2 (New England Biolabs). All PCRs were performed using Q5® High-Fidelity 2X Master Mix (NEB). Pooled samples were sequenced on a Miseq (Illumina), using 50-nt reads and collecting approximately 500-1000 reads per gRNA in the library.

The library prep primers were as follows:

PCR1:

U6_Bc_r1seq_halftail (24 distinct versions of this primer with staggered-length in-line barcodes denoted here as NNNNN were used)

5’ CTTTCCCTACACGACGCTCTTCCGATCT NNNNN GGAAAGGACGAAACACCG 3’

gRNAFE_r2seq_halftail

5’ GACTGGAGTTCAGACGTGTGCTCTTCCGATCTGCCTTATTTAAACTTGCTATGCTGT 3’

PCR2:

r1seq_fulltail

5’ AATGATACGGCGACCACCGAGATCTACACTCTTTCCCTACACGACGCTCTTC 3’

r2seq_fulltail (up to 8 distinct indexed versions of this primer were used to maximize pooling) 5’ CAAGCAGAAGACGGCATACGAGATNNNNNNNNGTGACTGGAGTTCAGACGTGTGCT 3’

CRISPR-Cas9 screening analysis was performed using MAGeCK RRA paired analysis comparing bottom (treatment) and top (control) sorted bins using non-targeting gRNAs as controls.

### Individual gRNA cloning

Oligonucleotides were ordered in the following format: GGAAAGGACGAAACACCG [protospacer] GTTTAAGAGCTATGCTGGAAAC and directly cloned into a lentiviral vector with a gRNA expression cassette, TagBFP fluorescence, and Puromycin resistance (addgene cat#230935) using BsmbI digest and NEBuilder HiFi DNA Assembly (NEB, E2621S). NEB Stable Competent E. coli cells (NEB, C3040H) were transformed using the assembly product, cultured overnight at 30C, and mini-prepped using the Qiagen Spin Miniprep Kit (Qiagen, 27104).

### Lentiviral packaging

HEK293T cells were seeded at 1.37 x 105 cells/cm^2^ in a 12 well plate (individual gRNAs) or 10-cm plate (gRNA libraries) in complete OptiMEM. Cells were transfected with psPAX2 (Addgene, 12260), pMD.2G (Addgene, 12259) and the respective transfer plasmid using Lipofectamine 3000 Transfection Reagent (ThermoFisher, L3000015). Lentivirus was collected after 24, 48, and 72 (for gRNA libraries) hours, filtered using a 0.45 µM filter and concentrated 50X using Lenti-X Concentrator (Takara, 631232).

### Individual gRNA RT-qPCR validation

i^3^N iPS dCas9^KRAB^ cells were seeded and transduced at 1.5 x 10^4^ cells/cm^2^ on a 24 well plate in 3-6 biological replicates per gRNA. Day 0 iNeurons were seeded and transduced at a density of 2 x 10^5^ cells/cm^2^ on a 6 well plate in 3-6 biological replicates. All cells were transduced with lentivirus at high MOI. iPSCs were selected with 0.5 µg/mL Puromycin on days 2-7. On Day 7, RNA was harvested from the cells using Qiagen RNeasy Mini kit (Qiagen, 74104) and DNase treated using RQ1 RNase free DNase (Promega, M6101). cDNA was generated using ProtoScript First Strand cDNA Synthesis Kit (NEB, E6300S) and a poly-T primer. RT-qPCR was performed using SensiMix SYBR Master Mix (OriGene, QP100001) using the following primers and cycling conditions:

SV2A_FW: CGCCTTTCCTTCTGTGTTTGCC

SV2A_RV: GAGAACACTCGCTCAGGATGTC

FAM120A_FW: CCGTCTGCATGGCCCACTG

FAM120A_RV: AGAGTCATACGCAACCAAGCCA

GAPDH_FW: GTCTCCTCTGACTTCAACAGCG

GAPDH_RV: ACCACCCTGTTGCTGTAGCCAA

1-2 µL template cDNA

1X SensiMix SYBR 2X Master Mix

0.5 µM Fw primer

0.5 µM Rv primer

dH2O to total volume of 15 µl

95C | 95C 60C 72C | 72C

10 min |15 sec 15 sec 15 sec | 5min 40 cycles

Fold-change in *FAM120A and SV2A* expression was calculated using the ΔΔCt method and normalized to a non-targeting control gRNA.

### GWAS loci and RE overlap with eHi-C loops

iPSC, iNeuron, fetal cortex, and adult cortex eHi-C data were obtained from NCBI GEO accession number GSE115407 (hg19). To annotate TSSs in the human genome (hg19), we used the TxDb.Hsapiens.UCSC.hg19.knownGene transcript annotation package from Bioconductor. TSSs were extracted by identifying the start position of each transcript, accounting for strand orientation. Specifically, the promoters() function from the GenomicRanges package was used with parameters upstream = 5000 and downstream = 5000 to define a 10 kb window at each TSS. To identify putative promoter-loop connections to SCZ-associated GWAS loci, we overlapped loop anchors in GWAS loci with the TSS coordinates described above using the findOverlaps() function. The resulting overlaps represent genomic regions where SCZ GWAS regions contact SCZ WES gene promoters.

To map CRISPRi-identified RE-promoter interactions for the three WES genes (*FAM120A*, *SV2A*, and *STAG1*), the above list of eHi-C promoter-GWAS region loops was overlapped with hg19 RE coordinates. We identified hg19 TSSs for the three WES genes and defined ±5 kb promoter windows. eHi-C anchors overlapping these promoter windows were removed so that only loops in which the other anchor contained a RE were retained.

### TFBS motif disruption

To predict transcription factor binding sites disrupted by variants, we used motifbreakR. Variants were formatted as GRanges objects containing chromosome, position (hg19), reference and alternate alleles, and unique SNP identifiers. We assessed TF motif disruption using the JASPAR human motif collection. P-values were then computed using calculatePvalue() with a granularity of 1×10⁻⁶, and variant-specific motif effects were visualized using plotMB.

### C9orf129 overexpression

i^3^N iPS dCas9^KRAB^ cells were seeded at a density of 4.2 x 10^4^ cells/cm^2^ on a 10-cm plate and were transduced with sgControl or sg*C9orf129*. On Day 3, the cells were re-seeded at 4.2 x 10^4^ cells/cm^2^ on a 6-well plate in 3-6 biological replicates and were transduced with either a scrambled control or *C9orf129* overexpression vector. The cells were selected with 0.5 µg/mL puromycin on days 1-7 and 2.5 μg/mL blasticidin on days 4-7. On Day 7, RT-qPCR and HCR-flowFISH were performed as described previously.

### Activity normalized prime editing library cloning

All oligonucleotide libraries (**Tables S12-16**) were ordered in the following sequence format:

5’ GTGGAAAGGACGAAACACC [19-nt protospacer] GTTTCAGAGCTATGCTGGAAACAGCATAGCAAGTTGAAATAAGGCTAGTCCGTTATCAACTTGAAAAAGTGGCACCGAGTCGGTGC [PBS/RT] CGCGGTTCTATCTAGTTACGCGTTAAACCAACTAGAATTTTTT [reporter] [7-nt barcode] AGATCGGAAGAGCACACGTCT 3’

pegRNA libraries were amplified by PCR using Q5® High-Fidelity 2X Master Mix (NEB) using the following primers and annealing temperatures:

pegRNA_60bp_fw: 5’

TAACTTGAAAGTATTTCGATTTCTTGGCTTTATATATCTTGTGGAAAGGACGAAACACCG 3’

pegRNA_r2seq_i721_BsrGrv: 5’

TTTAAAACTTTATCCATCTTTGCATGTACAGAAGACGGCATACGAGATNNCTGCGCGTGACTGGAGTTCAGA CGTGTGCTCTTCCGATCT3’

Ta = 69 °C

pegRNA libraries were then cloned into a lentivirus activity-normalized U6 promoter IN-PE2-SSB expression vector with puromycin resistance (pLenti-AN-U6-IN-PE2-SSB-puroR) through BsmBI vector digest and NEBuilder HiFi DNA assembly. Assembly reactions were electroporated to Lucigen Endura electrocompetent E. coli (60242-1), ensuring >100-fold representation of each pegRNA.

### Region #1 and region #2 prime editing timecourse

Two libraries installing SNPs or 5 bp deletions at a highly accessible (region #1) or lowly accessible (region #2) ATAC peak were cloned according to the activity normalized prime editing library cloning protocol outlined above. i^3^N iPSCs were seeded at a density of 4.17×10^4^ cells/cm^2^ on a 6-well plate in three biological replicates in 10 µM Y-27632. The cells were transduced with region #1 or region #2 pegRNA libraries at an MOI of 0.3 as determined by titration. One day post-transduction, 10 μM Y-27632 was removed. Two days post-transduction, cells were treated with 0.5 µg/mL of puromycin for the remainder of the experiment and with or without 1 mM VPA. 14 days post-transduction, each cell population was split into two conditions and seeded at a density of 2.08×10^4^ cells/cm^2^ on a 6-well plate in the presence of 10 μM Y-27632, with or without transduction of hMLH1dn-BlastR lentivirus. One day post-hMLH1dn-BlastR transduction, 10 μM Y-27632 was removed. Two days post-transduction, cells transduced with hMLH1dn-BlastR were treated with 2.5 µg/mL blasticidin. Days 7, 14, 21, and 28 post-library transduction, a portion of cells from non-hMLH1dn-BlastR transduced conditions were used as input for gDNA isolation using an Invitrogen PureLink Genomic DNA Mini Kit, with hMLH1dn-BlastR transduced conditions collected at Day 28 for gDNA isolation. The endogenous and reporter regions were amplified, barcoded, and sequenced as outlined below.

### NGS library preparation of endogenous sites

The 5’ to 3’ regions flanking endogenous pegRNA edit sites were amplified from gDNA using distinct in-line barcodes. PCR2 was performed to add full-length Illumina sequencing adapters using internally ordered primers with equivalent sequences to NEBNext Index Primer Sets 1 and 2. All PCRs were performed using Q5® High-Fidelity 2X Master Mix (NEB). Pooled samples were sequenced on a Miseq or MiSeq i100 (Illumina), using 150-nt paired or single-end reads and collecting approximately 40,000-150,000 reads per sample.

The library prep primers were as follows:

PCR1:

Bc_r1seq_halftail (distinct versions of this primer with staggered-length in-line barcodes denoted as NNNNN were used)

5’ CTTTCCCTACACGACGCTCTTCCGATCT NNNNN [Region-specific fw primer] 3’ r2seq_halftail

3’ GACTGGAGTTCAGACGTGTGCTCTTCCGATCT NNN [Region-specific rv primer] 5’

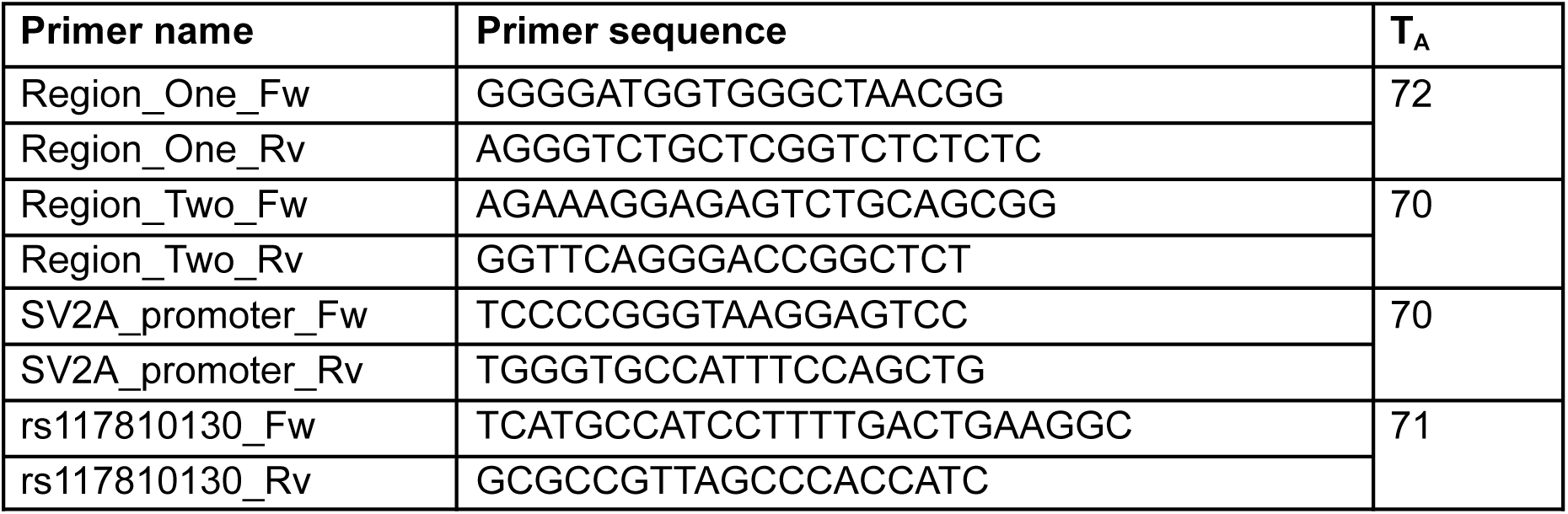

PCR2:

r1seq_fulltail

3’ AATGATACGGCGACCACCGAGATCTACACT CTTTCCCTACACGACGCTCTTC 5’

r2seq_fulltail (up to 8 distinct indexed versions of this primer were used to maximize pooling)

5’ CAAGCAGAAGACGGCATACGAGAT NNNNNNNN GTGACTGGAGTTCAGACGTGTGCT 3’

Pooled samples were sequenced on a Miseq or MiSeq i100 (Illumina), using 150-nt paired-end reads and collecting approximately 100,000-150,000 reads per sample. For the two region timecourse, reads were aligned to the reference sequences using CRISPresso2.

The linear regression model was calculated by fitting a line to the scatter plot of log_10_(50 bp window average ATAC-score + 1) vs. endogenous/reporter editing at week 4 with VPA and MMRi treatment.

### NGS library preparation of pegRNA and reporter

gDNA was isolated from cells and used to amplify the U6-3’ site to the region downstream of the pegRNA barcode with different in-line barcodes. PCR2 was performed to add full-length Illumina sequencing adapters and i5 indices. All PCRs were performed using Q5® High-Fidelity 2X Master Mix (NEB).

The library prep primers were as follows:

PCR1:

U6_PE1_BC (Up to 12 distinct versions, with up to 9-nt barcode, indicated as NNNNNNNNN) 5’ CTTTCCCTACACGACGCTCTTCCGATCT NNNNNNNNN GGAAAGGACGAAACACCG 3’

P7_anchor

3’ CAAG CAGAAGACGGCATACGAGAT 5’

PCR2:

NEBNext_i5 (8 distinct versions with unique 8-nt barcodes, indicated by NNNNNNNN)

5’ AATGATACGGCGACCACCGAGATCTACAC NNNNNNNN

ACACTCTTTCCCTACACGACGCTCTTCCGATC*T 3’

P7_anchor:

3’ CAAG CAGAAGACGGCATACGAGAT 5’

Pooled samples were sequenced using MiSeq i100 (Illumina), using 150-nt paired-end reads and collecting ∼1,000 reads per library member.

### Prime-editing variant screen design

The All dbSNP(155) database containing all Short Genetic Variants from dbSNP Release 155 was queried for variants in each of the three expanded GWA loci containing the expression ‘common’ in the ucscNotes column. The list was further filtered for variants meeting at least one of the following criteria 1) variants that map within CRISPRi-identified REs, 2) common variants in UTRs or splice sites of the WES gene, or 3) SCZ FINEMAP variants. REF and ALT variants were identified based on genotyping data described in the “Whole genome sequencing and variant calling” methods section. Additionally, synthetic 5 bp deletion variants centered on any SNPs and 10 synthetic 5 bp deletion sequences within ±100 bp of the WES gene TSS were added to the list. pegRNAs were generated using PRIDICT2.0. pegRNAs mutating the PAM or gRNA seed region were removed. pegRNAs installing variants that were not in **Table S11** and did not have at least one pegRNA with an MMR-proficient prime editing efficiency score above 18 from PRIDICT2.0 were removed, as well as their corresponding pegRNAs installing synthetic 5 bp deletion sequences. pegRNA libraries were formatted and cloned as described in the “Activity normalized prime editing library cloning” section above.

### PE HCR-FlowFISH screens

WTC-11 cells were seeded at a density of 2.11×10^4^ cells/cm^2^ on 15-cm plates in four biological replicates in the presence of 10 µM Y-27632. The cells were transduced with prime editing library lentivirus at a multiplicity of infection (MOI) of 0.3 as determined by titration. Two days post-transduction, cells were treated with 1 mM VPA for two days and 0.5 μg/mL puromycin for the remainder of the screen. On Day 7, cells were seeded at a density of 1.28×10^4^ cells/cm^2^ on 15-cm plates and transduced with hMLH1dn-BlastR at high MOI in the presence of 10 μM Y-27632. Y-27632 was removed after 24 hours. Two days post-transduction, cells were treated with 1 μg/mL blasticidin for the remainder of the screen. Three weeks post-library transduction, cells were lifted off the plate using accutase and used as input for genomic DNA (gDNA) isolation or the HCR-flowFISH protocol detailed above.

Cells were sorted into two bins (bottom 20% and top 20%) using a Sony Sorter flow cytometer. After sorting, gDNA was isolated from cells and used as input for NGS library preparation of pegRNA and reporter regions as detailed above.

### PE HCR-FlowFISH screen analysis

Reporter editing was calculated using the average reporter editing observed in bulk unsorted replicates. We input pegRNA counts into BEAN^51^ to calculate effect sizes. BEAN was run with default parameters, pegRNAs grouped by edit, and scaled by ATAC score (minimum scaling of 0.1). Matched control pegRNAs were scaled equally to the corresponding targeting pegRNA. To calculate variant effect sizes and z-scores relative to matched controls, we used the following equation: Effect size = ALT - REF; Z-score = (ALT - REF) / √(σ^2^_ALT_ + σ^2^_REF_).

### Individual pegRNA cloning

gBlocks containing pegRNA sequences were ordered in the following sequence format:

TAACTTGAAAGTATTTCGATTTCTTGGCTTTATATATCTTGTGGAAAGGACGAAACACCG [19-nt protospacer] GTTTCAGAGCTATGCTGGAAACAGCATAGCAAGTTGAAATAAGGCTAGTCCGTT ATCAACTTGAAAAAGTGGCACCGAGTCGGTGC [RT] [PBS] CGCGGTTCTATCTAGTTACGCGTTAAACCAACTAGAA TTTTTTT [reporter] [7-nt barcode] AGATCGGAAGAGCACACGTCTGAACTCCAGTCACGCGCAGAGATCTCGTATGCCGTCTTCTGTACATGCAAAGATGGATAAAGTTTTAAA

gBlocks were inserted into pLenti-AN-U6-IN-PE2-SSB-puroR through BsmBI vector digest and NEBuilder HiFi DNA assembly. NEB Stable chemically competent E. coli cells (NEB, C3040H) were transformed using the assembled product, cultured overnight at 30C, and prepped using a QIAprep Spin Miniprep Kit (Qiagen, 27104).

### Clonal isolation

Two hours prior to transfection, i^3^N iPSCs were seeded at a density of 1.04×10^5^ cells/cm^2^ on a matrigel-coated 12-well plate in 10 μM Y-27632. Cells were transfected using Lipofectamine (ThermoFisher, L3000008) with individual pegRNA constructs (1 μg DNA, 2 μL p3000, 3 μL Lipofectamine 3000 per well). Transfected cells were selected with 1.0 μg/mL puromycin for 48 hours beginning one day post-transfection. Cells were transfected up to three times with one week between transfections. 1.5-2 weeks after the final transfection, cells were isolated by FACS sorting onto 96-well plates in the presence of 10 µM penicillin-streptomycin and 10 μM Y-27632. For the three days following, half of the media (50 μL) was removed from each well and replaced with 50 μL of media without penicillin-streptomycin and Y-27632. Clonal populations were expanded to 24- and 12-well plates and gDNA and RNA were isolated, respectively, using the Invitrogen PureLink Genomic DNA Mini Kit (ThermoFisher, K182002) and Qiagen RNeasy Mini kit (Qiagen, 74134), respectively. gDNA was used as input for endogenous site library prep, while RNA was used as input for cDNA generation and RT-qPCR as described above.

Sequencing reads were trimmed to a 51-bp window centered on the edit using fastx_trimmer (FASTX Toolkit v0.0.14). We used CRISPResso2 (v2.2.3, minimum bp quality of 30 or change to N, minimum average read quality of 30) to align the trimmed FASTQ files to the reference sequence. Genotypes were assigned based on the following thresholds: REF/REF (REF: 80-100%; ALT: 0-20%), REF/ALT (REF: 30-70%; ALT: 30-70%), and ALT/ALT (REF: 0-20%; ALT: 80-100%).

To differentiate clonal lines into iNeurons, cells were seeded at 9.1×10^4^ cells/cm^2^ on a matrigel-coated 24-well plate in 10 μM Y-27632. The following day (day −3), media was replaced with pre-differentiation medium (PDM; Knockout DMEM/F-12 (1X) supplemented with N-2 (1X), NEAA (1X), BDNF (10ng/mL), NT-3 (10 ng/mL), laminin (1 µg/mL), and doxycycline (2 µg/mL)). Media was replaced with fresh PDM for the next two days (days −2 and −1). On day 0, PLO-coated 12-well plates were washed three times with PBS and seeded with maturation medium (Neurobasal-A (0.5X), DMEM/F12-HEPES (0.5X), NEAA (1X), B-27 (0.5X), N-2 (0.5X), GlutaMAX (0.5X), BDNF (10 ng/mL), NT-3 (10 ng/mL), laminin (1 µg/mL), and doxycycline (2 µg/mL)). Cells were lifted with accutase, counted, and replated in maturation medium at 1.04×10⁵ cells/cm^2^. On day 7, RNA was isolated using the Qiagen RNeasy Mini kit (Qiagen, 74134) and used as input for cDNA generation and RT-qPCR as described above.

### GFP reporter cloning

gBlocks were ordered in the following format: CTCACTCAGCCTGCATTTCTGCCAGGGCCCGCTCTAGACCTGCAGGAGGACCGGATCAACT [200 bp RE with ALT or REF variant] CATTGCGTGAACCGACACTAGAGGGTATATAATGGAAGCTCGACTTCCAGCTTGGCAATCCGGTACTGTGCAAAGTGAACACATCGCTAAGCGAAAGCTAAGACCGGTCGCCACCATGGTGAGCAAGGGCGAGGAGCTGT

TCA, resuspended to 5 ng/µL in 1X IDTE buffer, and directly cloned into a lentiviral GFP MPRA vector (Addgene plasmid #137725) using SbfI-HF and AgeI-HF digestion and NEBuilder HiFi DNA Assembly. NEB Stable Competent E. coli cells (NEB, C3040H) were transformed using the assembly product, cultured overnight at 30C, and mini-prepped using the Qiagen Spin Miniprep Kit (Qiagen, 27104).

### GFP reporter titration by qPCR

i^3^N iPSCs were seeded at a density of 7.81×10^3^ cells/cm^2^ on a 24-well plate in 10 µM Y-27632 and transduced with varying amounts of lentivirus in 2X increments. One day post-transduction, 10 µM Y-27632 was removed. Three days post-transduction the cells were used as input for gDNA isolation using an Invitrogen PureLink Genomic DNA Mini Kit and qPCR was performed to amplify viral DNA (WPRE.F and WPRE.R), plasmid backbone DNA (BB.F and BB.R) and gDNA (LP34.F and LP34.R) using the following primers and cycling conditions:

WPRE.F:: TACGCTGCTTTAATGCCTTTG

WPRE.R: GGGCCACAACTCCTCATAAAG

LP34.F: TCCTCCGGAGTTATTCTTGGCA

LP34.R: CCCCCCATCTGATCTGTTTCAC

BB.F: TGCCGCATAGTTAAGCCAGTA

BB.R: TCAAGCCTTGCCTTGTTGTAG

3.5 µL gDNA (10 ng/µL)

1X OneTaq® 2X Master Mix with Standard Buffer

0.5 µM Fw primer

0.5 µM Rv primer

0.75 µL EvaGreen^®^ Dye, 20X in Water dH2O to total volume of 15 µl

98C | 98C 50C 68C | 72C

10 min |15 sec 30 sec 60 sec | 5min 35 cycles

MOI was determined by calculating the relative amount of viral DNA over gDNA with the subtraction of the relative amount of backbone DNA.

### GFP reporter assay

For the iPSC GFP reporter assays, i^3^N iPSCs were seeded and transduced at 1.5×10^4^ cells/cm^2^ on matrigel-coated 24-well plates in 10 µM Y-27632 in 3-6 biological replicates. All cells were transduced with lentivirus at an MOI of 1.5. One day post-transduction, the media was replaced with fresh mTeSR, which was refreshed every 24 hours. Three days post-transduction, cells were lifted and analyzed for GFP expression using a Sony Sorter flow cytometer.

For the iNeuron GFP reporter assays, i^3^N iPSCs were seeded and transduced at a density of 1.5×10^4^ cells/cm^2^ on a matrigel-coated 24-well plate in 10 µM Y-27632 in three biological replicates. All cells were transduced with lentivirus at an MOI of 1.5. One day post-transduction, the media was replaced with fresh mTeSR, which was refreshed every 24 hours. Three days post-transduction, cells were replated at 1×10⁵ cells/cm^2^ onto matrigel-coated 24-well plates in mTeSR with 10 µM Y-27632. The following day (day −3), media was replaced with pre-differentiation medium (PDM; KnockOut DMEM/F-12 (1X) supplemented with N-2 (1X), NEAA (1X), BDNF (10 ng/mL), NT-3 (10 ng/mL), laminin (1 µg/mL), and doxycycline (2 µg/mL)) for three consecutive days. On day 0, PLO-coated 24-well plates were washed three times with PBS and seeded with maturation medium (Neurobasal-A (0.5X), DMEM/F12-HEPES (0.5X), NEAA (1X), B-27 (0.5X), N-2 (0.5X), GlutaMAX (0.5X), BDNF (10 ng/mL), NT-3 (10 ng/mL), laminin (1 µg/mL), and doxycycline (2 µg/mL)). Cells were lifted with accutase, counted, and replated in the maturation medium at 5×10^4^ cells/cm^2^. On day 7, iNeurons were dissociated using an accutase/papain solution at 20 units of papain per mL, quenched in papain quench solution, resuspended in FACS suspension media, and analyzed for GFP expression using a Sony Sorter flow cytometer.

### SV2A CRISPRa and MEA

i^3^N iPSC DHFR-dCas9-VPH cells were seeded at a density of 1.5×10^4^ cells/cm^2^ on a matrigel-coated 24-well plate in 10 µM Y-27632 and transduced with one of three CRISPRa *SV2A* gRNAs or one of two non-targeting controls in 3-4 biological replicates. The following day, the cell culture media was replaced without Y-27632. Two days post-transduction, cells were selected with 2 µg/mL puromycin. Following selection, cells were seeded at 1.5×10^4^ cells/cm^2^ on a matrigel-coated 24-well plate in 10 µM Y-27632. The following day (day −3), media was replaced with pre-differentiation medium (PDM; KnockOut DMEM/F-12 (1X) supplemented with N-2 (1X), NEAA (1X), BDNF (10 ng/mL), NT-3 (10 ng/mL), laminin (1 µg/mL), and doxycycline (2 µg/mL)) for three consecutive days.

On day 0, cells were plated on a MEA 96-well plate (Axion Biosystems M768-tMEA-96W). To prepare the MEA plate, all wells were coated with fetal bovine serum for 60 minutes at 37C. Then, FBS was aspirated and the wells were washed three times with PBS. The wells were then coated with PDL for 60 minutes at 37C. Then, PDL was aspirated and the wells were washed three times with PBS. Next, the plate was left open in the biosafety cabinet to air dry for 60 minutes before plating cells. Cell were accutased and counted, then 5µL of pre-differentiated neurons at a concentration of 1×10^7^/mL resuspended in maturation medium (Neurobasal-A (0.5X), DMEM/F12-HEPES (0.5X), NEAA (1X), B-27 (0.5X), N-2 (0.5X), GlutaMAX (0.5X), BDNF (10 ng/mL), NT-3 (10 ng/mL), laminin (1 µg/mL), and doxycycline (2 µg/mL)) with TMP were plated at the center of each well covering the area of the well with the electrodes in up to four technical replicates per biological replicate. The plate was incubated at 37C for 45 minutes for the cells to attach, and 150 µL/well of maturation medium with TMP was gently added to each well. On day 7, half of the maturation medium was removed and the same amount was added back, without doxycycline, and with TMP. On day 14, half of the maturation medium was removed and twice the amount was added back, without doxycycline, and with TMP. MEA recording was taken at day 21 for viability and spontaneous activity on Axion Maestro Pro. Spike files were generated automatically by the software at the end of each recording and were used for downstream analysis.

On day 0, additional cells were plated for RT-qPCR analysis. PLO-coated 24-well plates were washed three times with PBS and seeded with maturation medium (Neurobasal-A (0.5X), DMEM/F12-HEPES (0.5X), NEAA (1X), B-27 (0.5X), N-2 (0.5X), GlutaMAX (0.5X), BDNF (10 ng/mL), NT-3 (10 ng/mL), laminin (1 µg/mL), and doxycycline (2 µg/mL)). Cells were seeded in the maturation medium at a density of 5×10^4^ cells/cm^2^. Cells were fed as described above at days 7 and 14. On days 7, 14, and 21 RNA was isolated using the Qiagen RNeasy Mini kit (Qiagen, 74134) and used as input for cDNA generation and RT-qPCR of *SV2A* as described above.

