## Supplemental Figures for "Interrogation of noncoding schizophrenia risk variants using CRISPR-based functional genomics"

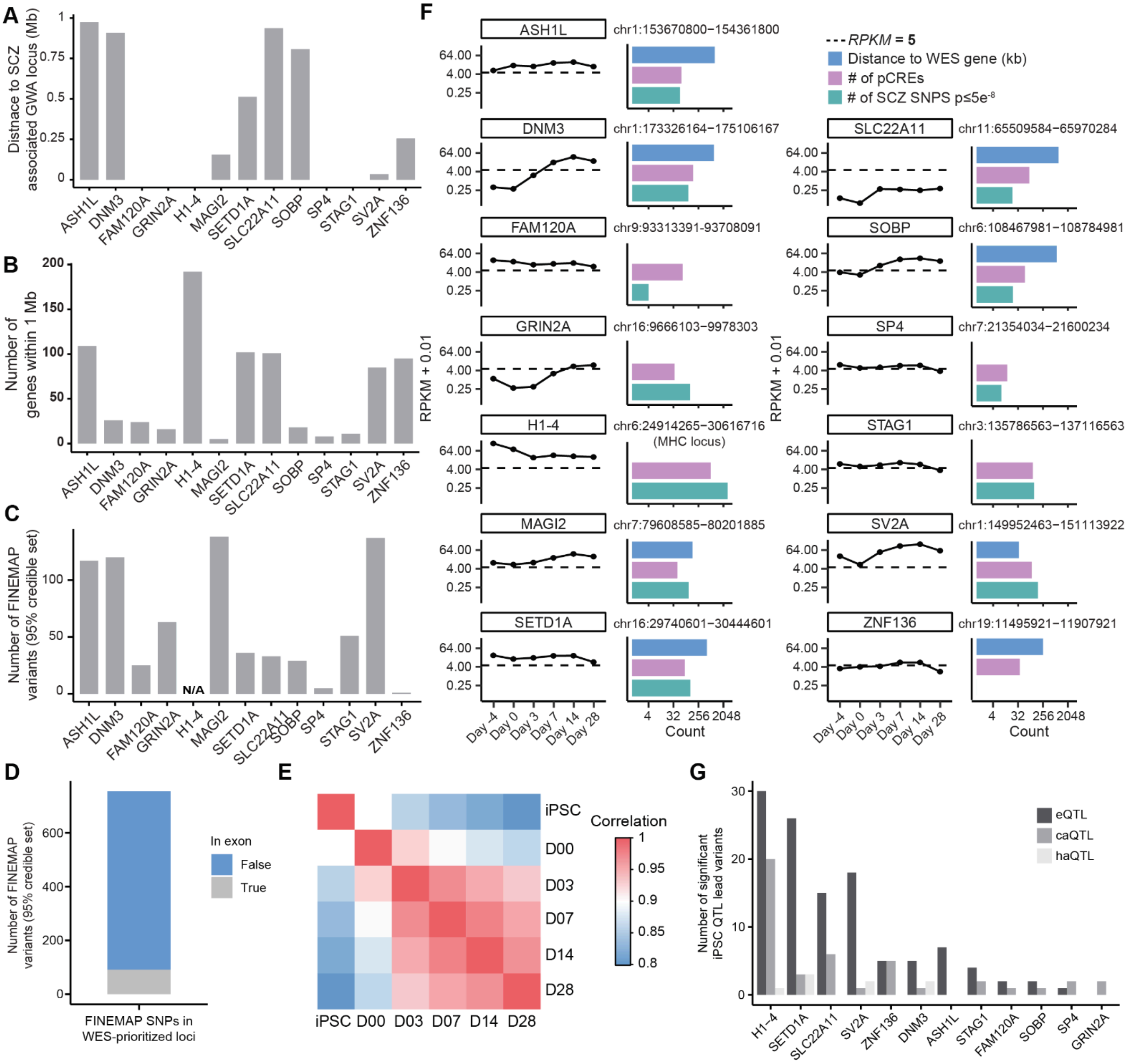


**Figure S1, Related to Figure 1.** A) Distance of 13 SCZ WES genes within 1 Mb of a SCZ GWA locus to the nearest SCZ GWA locus. B) Number of protein coding genes within 1 Mb of each of the 13 WES-prioritized GWA loci. Each locus is labeled with the corresponding putative SCZ gene target. C) Number of significant SNP associations within the 13 WES-prioritized GWA loci. Each locus is labeled with the corresponding putative SCZ gene target. D) Number of finemapped SNPs (95% credible set) in WES-prioritized GWA loci mapping within or outside of exonic regions. E) Spearman correlations of RNA-seq mean TPM per stage for 13,569 protein-coding genes detected in any stage. F) Left: Expression of 13 SCZ WES genes within 1Mb of a SCZ GWA locus in iPSCs or iNeurons throughout four weeks of differentiation. Right: Bar plot depicting the distance to the closest SCZ WES gene, number of pCREs, and number of SCZ SNPs with $P \leq 5e^{-8}$ for each of the 13 SCZ GWA loci. G) Number of iPSC xQTL lead variant signals in 13 WES-prioritized loci. Each locus is labeled with the corresponding putative SCZ gene target.

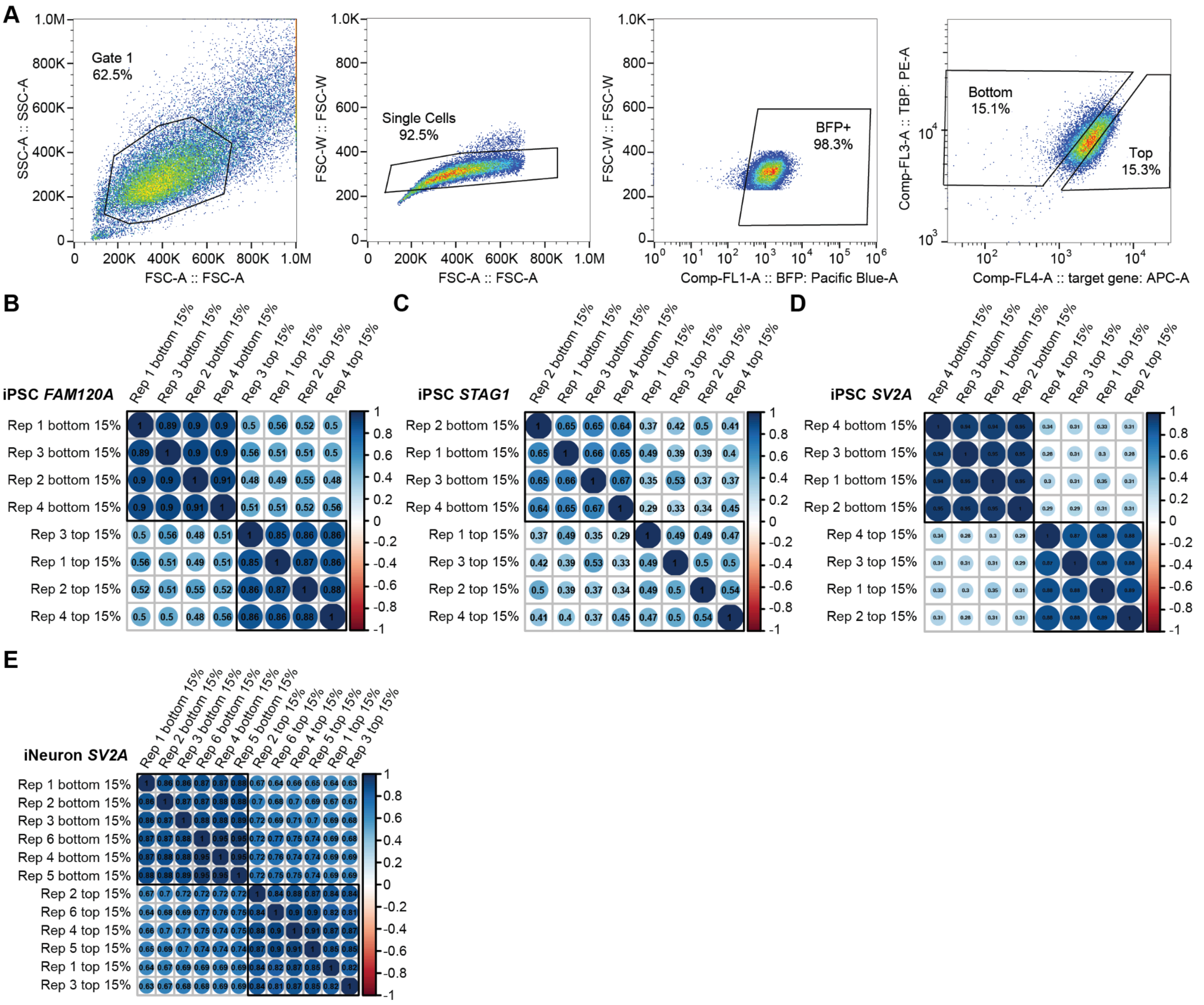


**Figure S2, Related to Figure 2.** A) Representative FACS plots for sorting i$^3$N iPSCs cells into bottom 15% and top 15% fluorescent bins after HCR-flowFISH for *FAM120A* expression, normalized to *TBP.* Pearson correlation of the reads per million (RPM) of the bottom 15% and top 15% bins for the (B-D) iPSC CRISPRi screens or (E) iNeuron CRISPRi screen.

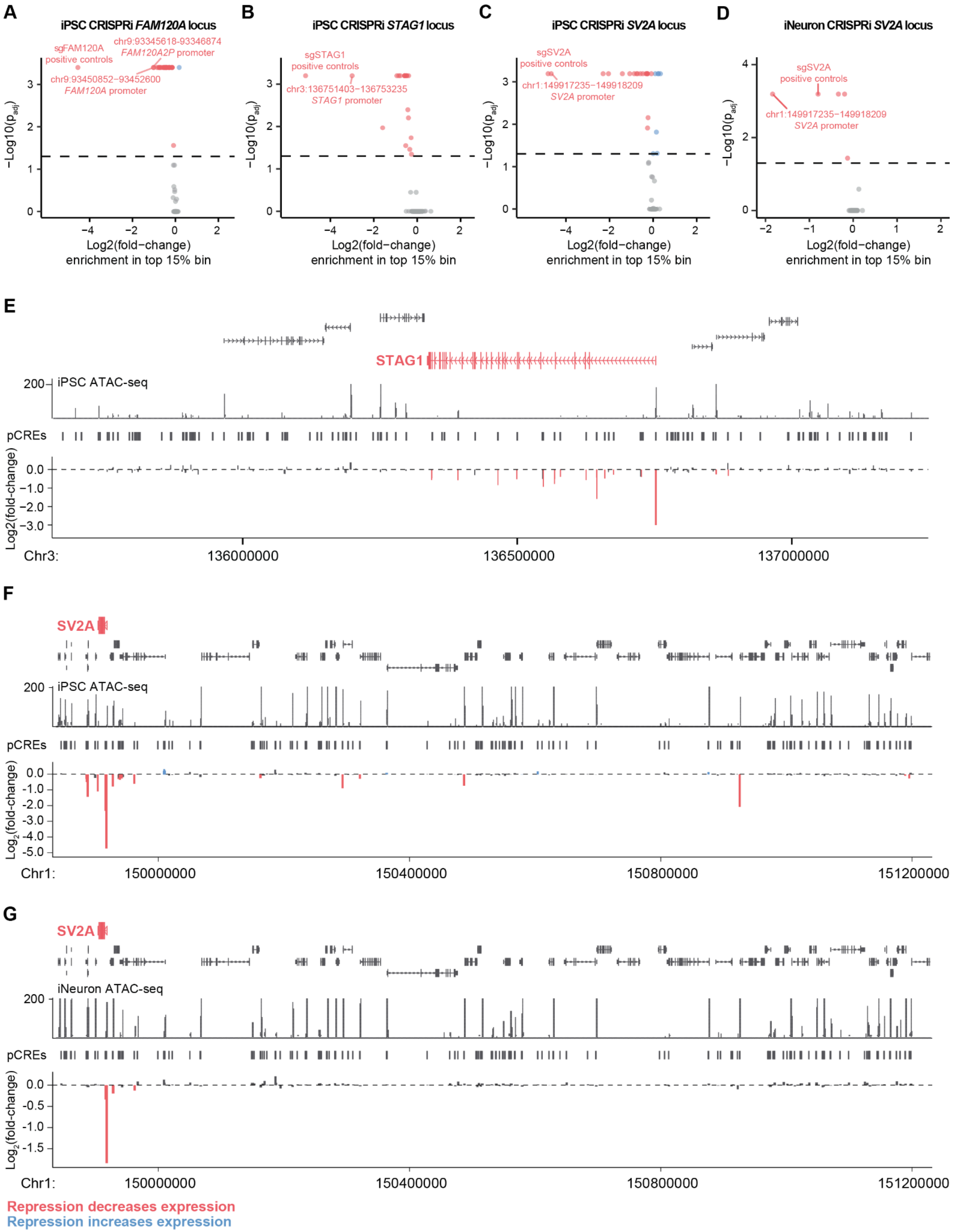
A
iPSC CRISPRi FAM120A locus
sgFAM120A
positive controls
chr9:93345618-93346874
FAM120A2P promoter
chr9:93450852-93452600
FAM120A promoter
B
iPSC CRISPRi STAG1 locus
sgSTAG1
positive controls
chr3:136751403-136753235
STAG1 promoter
C
iPSC CRISPRi SV2A locus
sgSV2A
positive controls
chr1:149917235-149918209
SV2A promoter
D
iNeuron CRISPRi SV2A locus
sgSV2A
positive controls
chr1:149917235-149918209
SV2A promoter
−Log10(p_adj)
Log2(fold−change)
enrichment in top 15% bin
E
STAG1
iPSC ATAC-seq
pCREs
Log2(fold-change)
Chr3:
136000000
136500000
137000000
F
SV2A
iPSC ATAC-seq
pCREs
Log2(fold-change)
Chr1:
150000000
150400000
150800000
151200000
G
SV2A
iNeuron ATAC-seq
pCREs
Log2(fold-change)
Chr1:
150000000
150400000
150800000
151200000
Repression decreases expression
Repression increases expression

**Figure S3, Related to Figure 2.** Volcano plots showing the $\log_2$(fold-change) enrichment of pREs in the top vs bottom CRISPRi HCR-flowFISH bins and their corresponding $-\log_{10}$(p-values) in $i^3$N iPSCs for pREs in SCZ-associated GWA loci prioritized by (A) *FAM120A,* (B) *STAG1*, or (C) *SV2A*. REs significantly decreasing expression of the target gene are depicted in red. REs significantly increasing expression of the target gene are depicted in blue. D) Volcano plot showing the $\log_2$(fold-change) enrichment of pREs in the top vs bottom CRISPRi HCR-flowFISH bins and their corresponding $-\log_{10}$(p-values) in iNeuron cells for pREs in SCZ-associated GWA loci prioritized by *SV2A*. Locus view displaying the (E) *STAG1* or (F) *SV2A* GWA loci. From top to bottom: GENCODE V48 genes and predicted genes; $i^3$N iPSC ATAC-seq scores; pREs identified through called ATAC-seq peaks; pRE level $\log_2$(fold-change) in enrichment in top vs bottom 15% bins after HCR-flowFISH and FACS sorting of $i^3$N iPSCs transduced with the pRE targeting gRNA libraries. G) Locus view displaying the *SV2A* GWA locus. From top to bottom: GENCODE V48 genes and predicted genes; iNeuron ATAC-seq scores; pREs identified through called ATAC-seq peaks; pRE level $\log_2$(fold-change) in enrichment in top vs bottom 15% bins after HCR-flowFISH and FACS sorting of iNeurons cells transduced with the pRE targeting gRNA libraries.

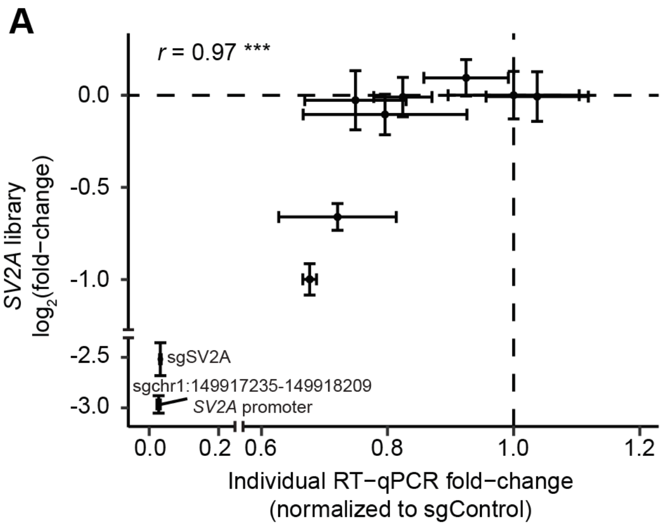


**Figure S4, Related to Figure 2.** A) Pearson correlation between expression of *SV2A* in an individual gRNA validation using RT-qPCR and the *SV2A* pRE library gRNA $\log_2$(fold-change) in HCR-flowFISH screening of iNeurons. (∗p < 0.05, ∗∗p ≤ 0.01, ∗∗∗p ≤ 0.001).

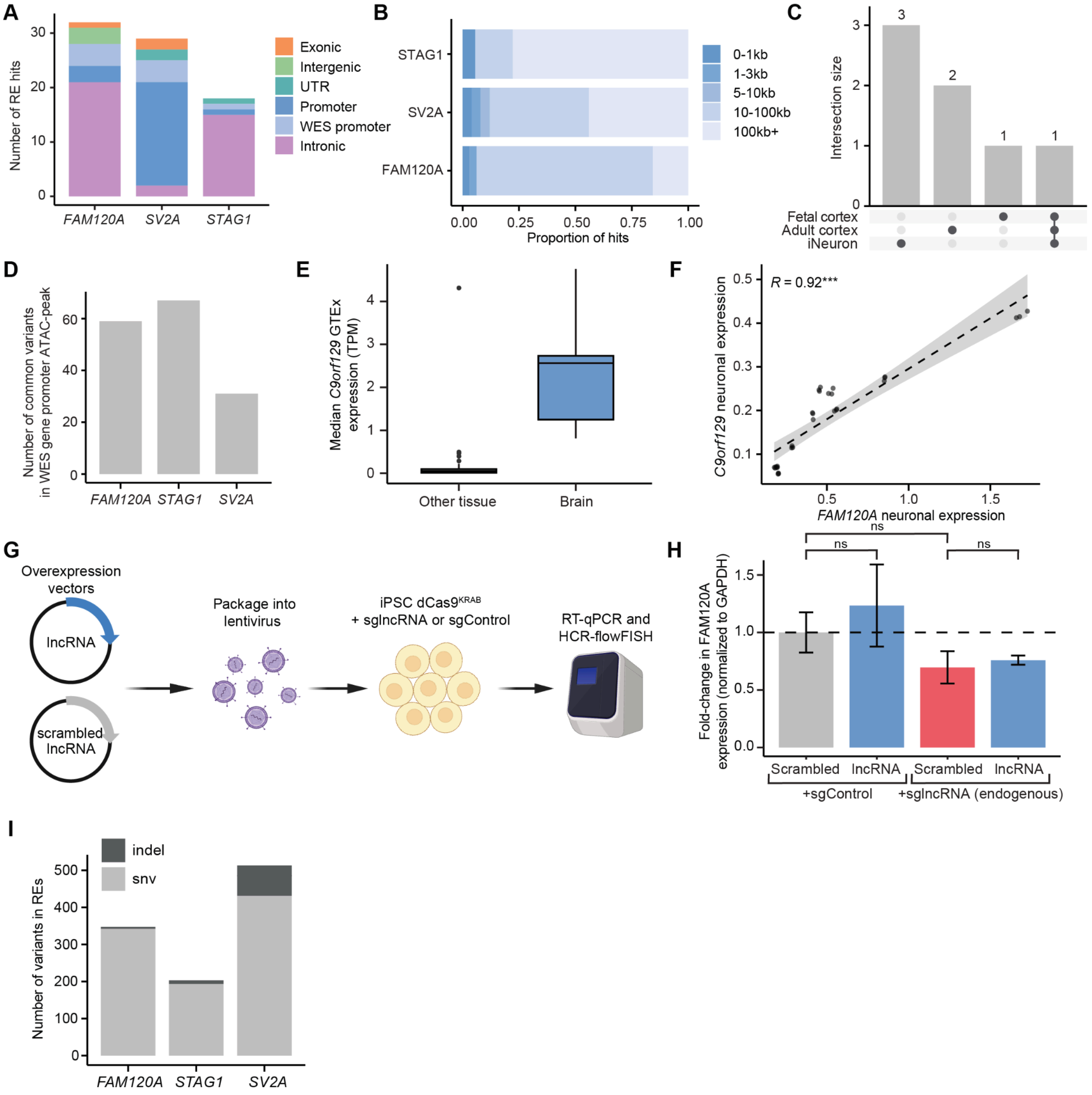


**Figure S5, Related to Figure 3.** A) Genomic annotations of RE hits, categorized as intronic, promoter, UTR, intergenic, or exonic. B) Distance of RE hits from the transcription start site (TSS) of the indicated gene target, binned as 0-1 kb, 1-3 kb, 5-10 kb, 10-100 kb, and 100 kb+. C) UpSet plot showing intersections of REs and chromatin loops identified in fetal cortex, adult cortex, and iNeuron Hi-C datasets[1]. D) Number of common variants mapping in the ATAC-peak at the promoter of FAM120A, STAG1, or SV2A. E) GTEx median TPM expression of *FAM120A* in the brain and all other GTEx tissues. G) Correlation of *FAM120A* and *C9orf129* expression in primary neurons from a single-cell atlas of the developing brain[2]. H) Schematic depicting the experimental workflow for overexpression of *C9orf129*. Overexpression vectors for *C9orf129* (lncRNA) or a scrambled negative control were packaged into lentivirus and delivered to $i^3N$ iPSC cells expressing sgC9orf129 or sgControl. *FAM120A* expression was analyzed using RT-qPCR. I) Normalized fold-change in *FAM120A* expression following overexpression of *C9orf129* lncRNA compared to scramble control in $i^3N$ iPSC expressing sgC9orf129 or sgControl as measured by RT-qPCR. Two-tailed t test. J) Number of common variants (SNVs and indels) located within CRISPRi-identified REs for *FAM120A*, *STAG1*, and *SV2A*. ($*p < 0.05$, $**p \le 0.01$, $***p \le 0.001$).

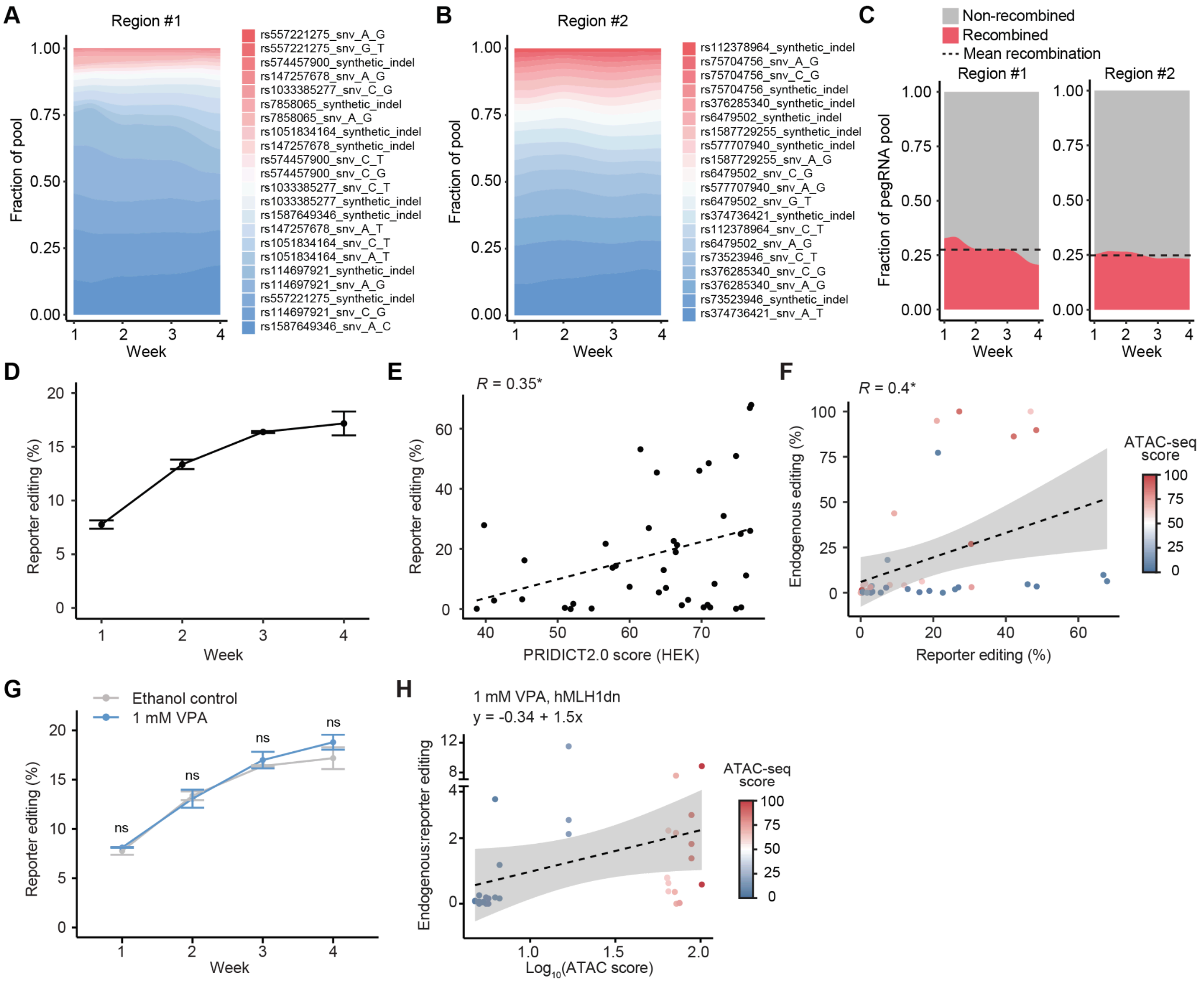


**Figure S6, Related to Figure 4.** Fraction of pegRNA-reporter NGS sequencing reads contributed by each pegRNA targeting (A) the highly accessible locus (region #1) or (B) the lowly accessible locus (region #2) throughout a four-week timecourse, colored by pegRNA identity. C) Proportion of recombined versus non-recombined pegRNA-reporter reads for both pegRNA libraries throughout a four-week time course. The dashed line indicates mean recombination frequency. D) Average reporter editing efficiency for all pegRNAs targeting region #1 and region #2 throughout a four-week time course. E) Pearson correlation between PRIDICT2.0 pegRNA efficiency scores (trained in HEK293T cells) and reporter editing in iPSCs for all pegRNAs, measured at four weeks post-transduction. F) Pearson correlation between endogenous and reporter editing efficiency for all pegRNAs targeting region #1 and #2, measured at four weeks post-transduction. Each pegRNA is colored based on its average ATAC-score within a 50-bp window of the targeted variant. G) Average reporter editing efficiency for all pegRNAs throughout a four-week time course with VPA or control treatment. Paired t test. H) Pearson correlation between endogenous:reporter editing ratio and chromatin accessibility (ATAC-seq score) following VPA and hMLH1dn treatment at four weeks post-transduction. Each pegRNA is colored based on its average ATAC-score within a 50-bp window of the targeted variant. ($*p < 0.05$, $**p \le 0.01$, $***p \le 0.001$).

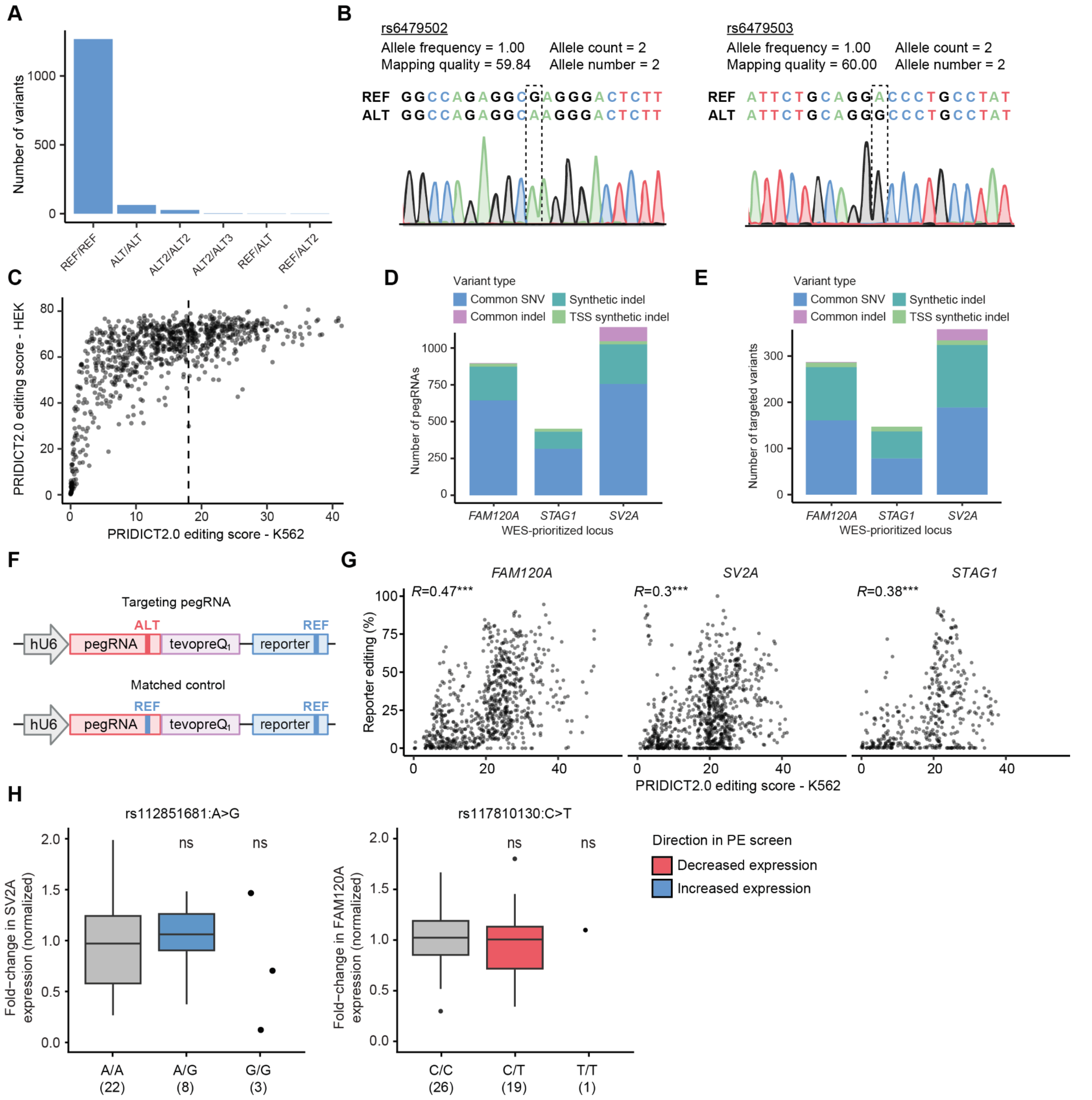


**Figure S7, Related to Figure 5.** A) Bar plot showing the number of i³N iPSC variants displaying different genotypes relative to the hg38 reference genome. B) Sanger sequencing of endogenous sites depicting two variants in which the i³N iPSCs harbor homozygous ALT variants relative to the hg38 reference genome. C) PRIDICT2.0 K562 and HEK editing scores for the top pegRNA for each variant (chosen based on K562 editing score). Bar plot depicting the number of D) pegRNAs per library and E) targeted variants per library. F) Schematic of targeting and matched control pegRNAs installing ALT or REF variants, respectively. G) Pearson correlation of reporter editing and PRIDICT2.0 scores for pegRNAs in the *FAM120A*, *SV2A*, or *STAG1* screens. H) Normalized *SV2A* or *FAM1120A* expression in iPSC edited clones across two variants and genotypes as measured by RT-qPCR. Number of clones in parentheses.Two-tailed t test. (*$p < 0.05$, **$p \le 0.01$, ***$p \le 0.001$).

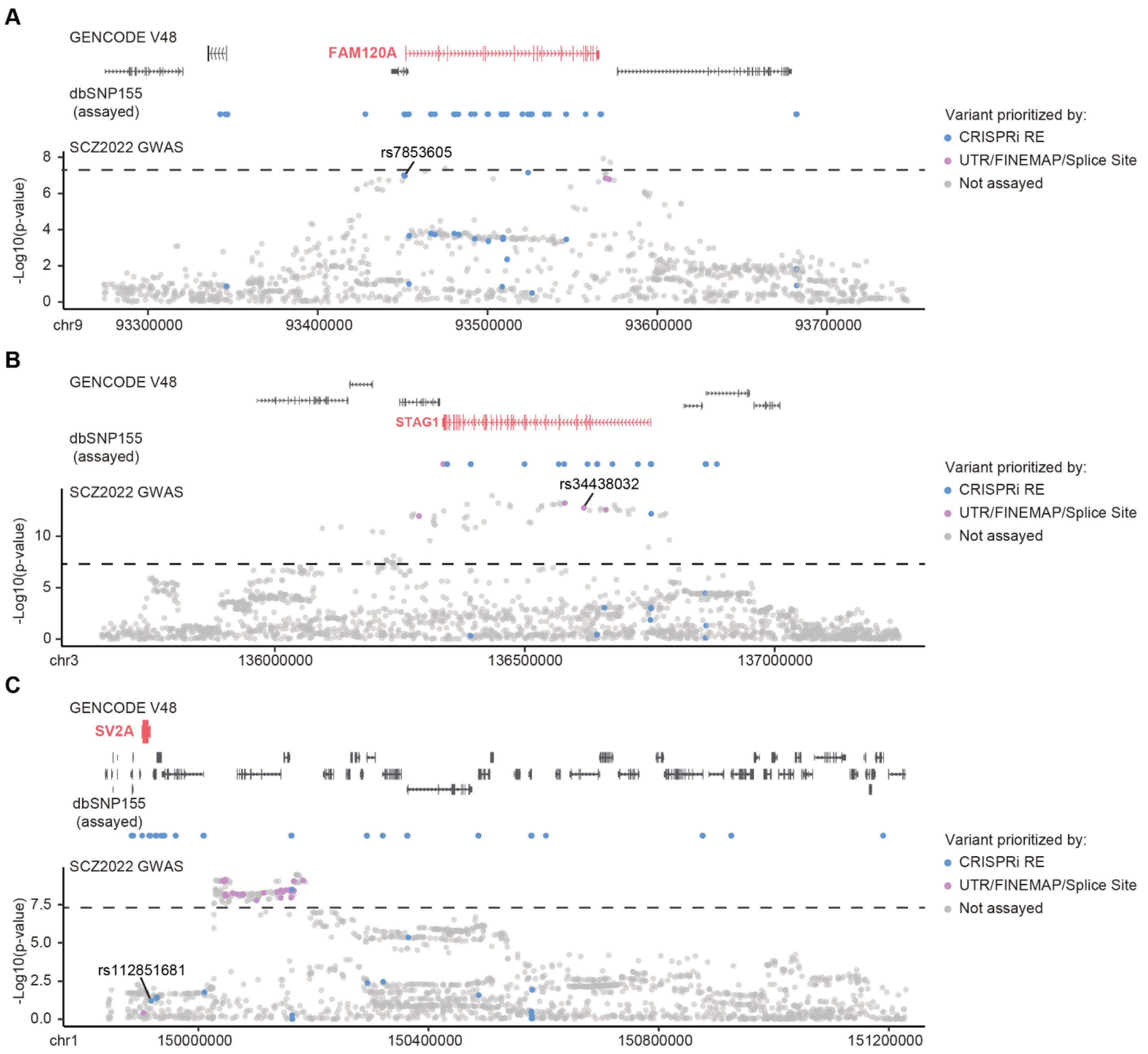


**Figure S8, Related to Figure 5.** Locus view displaying variants assayed via pooled prime editing screens in the A) *FAM120A*, B) *STAG1*, or C) *SV2A* GWA locus. From top to bottom: GENCODE V48 genes and predicted genes; dbSNP155 variants not in the SCZ2022 GWAS, but assayed in the pooled prime editing screens; dbSNP155 variants in the SCZ2022 GWAS, with variants assayed in the pooled prime editing screens highlighted in blue or purple.

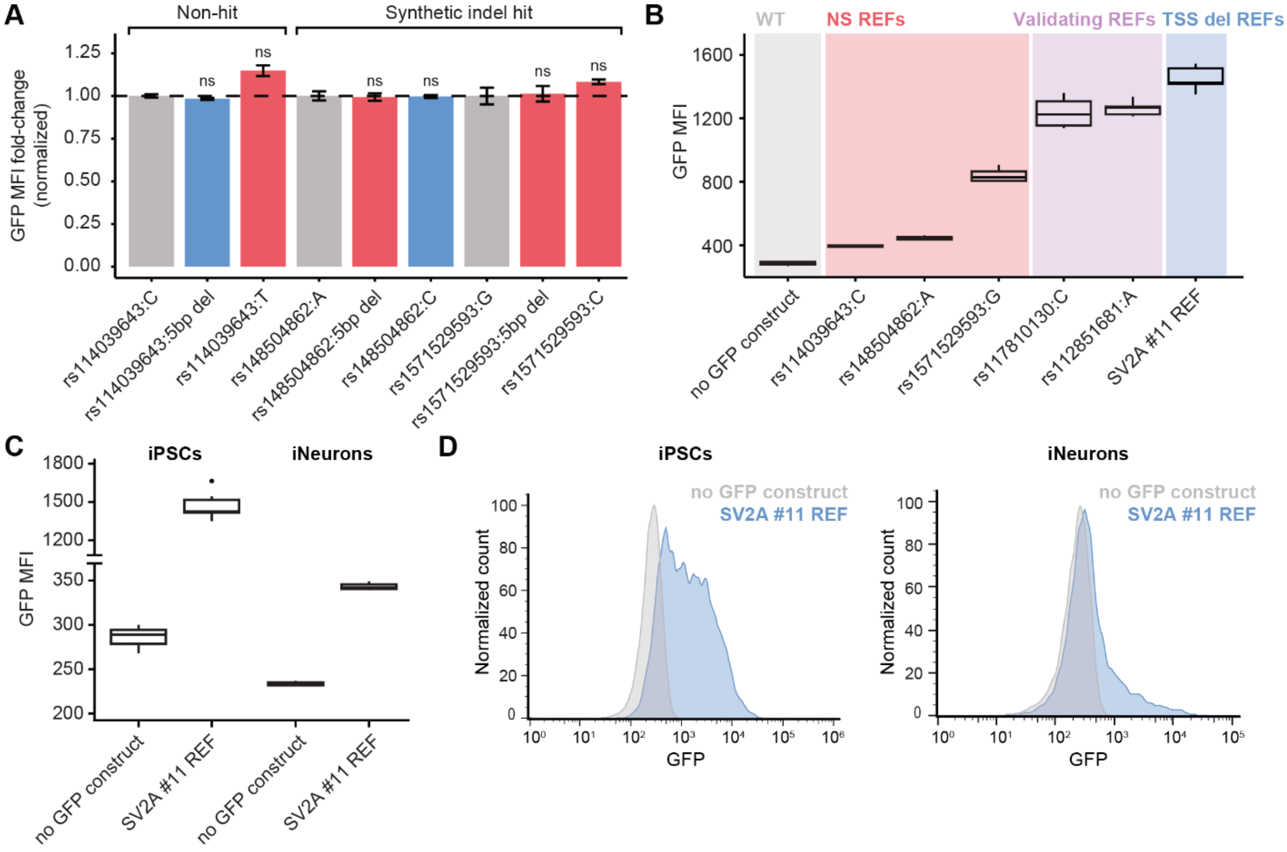


**Figure S9, Related to Figure 6.** A) Normalized fold-change in GFP MFI in iPSCs for three ALT 5 bp synthetic indels and their corresponding SNPs relative to the reference sequence. n=3-6. Two-tailed t test. B) GFP MFI in iPSCs for all tested reference sequences and no GFP (untransduced) cells. NS = not significant. C) GFP MFI in iPSCs and iNeurons for the 5 bp TSS deletion SV2A #11 reference sequence and no GFP (untransduced) cells. D) Representative GFP MFI histograms of iPSCs and iNeurons for the 5 bp TSS deletion SV2A #11 reference sequence and no GFP (untransduced) cells. (ns = not significant).

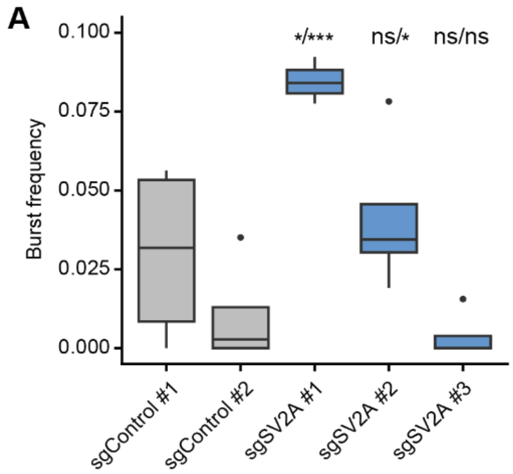

**Figure S10, Related to Figure 7.** A) Burst rate of D21 DHFR-dCas9-VPH iNeurons transduced with either non-targeting control or *SV2A* gRNAs. Two-tailed t test. (*$p < 0.05$, **$p \leq 0.01$, ***$p \leq 0.001$).
